# Heterogeneity of HCMV-Specific CD8^+^ T Cells is Established Early in Infection, Clonally Encoded, and Stable Over Time

**DOI:** 10.64898/2026.10.09.758110

**Authors:** Samuel D. Klebanoff, Matthew Lawrance, Kimberly O’Brien, Megan E. Smithmyer, Cate Speake, Amy S. Codd, Emily S. Ford, David M. Koelle, Jane H. Buckner, Evan W. Newell, Daniel J. Campbell

**Affiliations:** Benaroya Research Institute; University of Washington Department of Immunology; Fred Hutchinson Cancer Center; University of Washington Department of Medicine; University of Washington Department of Laboratory Medicine and Pathology; University of Washington Department of Global Health

## Abstract

Human cytomegalovirus (CMV) elicits lifelong memory T cell responses of striking size and phenotypic diversity, but how these diverse responses are established during acute infection and evolve over decades of chronic stimulation is poorly understood. We tracked CD8^+^ T cells specific for the viral antigens pp65 and IE1 by single-cell RNAseq, TCRseq, and CITEseq over a period of 16 months in a newly seroconverted donor, and in five latently infected donors over intervals ranging from 4.5 to 12.5 years. We found that individual T cell clones exhibited distinct, specificity-dependent transcriptional profiles, with pp65-specific cells skewed toward central memory phenotypes and IE1-specific cells enriched for effector and chronically activated states. Among pp65-specific T cells we identified a *CMC1*-expressing state not previously described in virus-specific T cells that shares transcriptional features with exhausted T cells in human tumors yet remains phenotypically stable and clonally intact for over a decade of viral latency. Distinct clonal phenotypes arose during acute infection and were remarkably stable during viral latency, and these phenotypes tracked with relative TCR avidity and TCR sequence. Our findings reveal that the phenotypic diversity of the human CMV-specific CD8+ T cell response is organized at the level of individual clones, with distinct functional identities established during acute infection and preserved with striking fidelity across decades of viral latency and chronic antigen exposure.

**Graphical Abstract:** 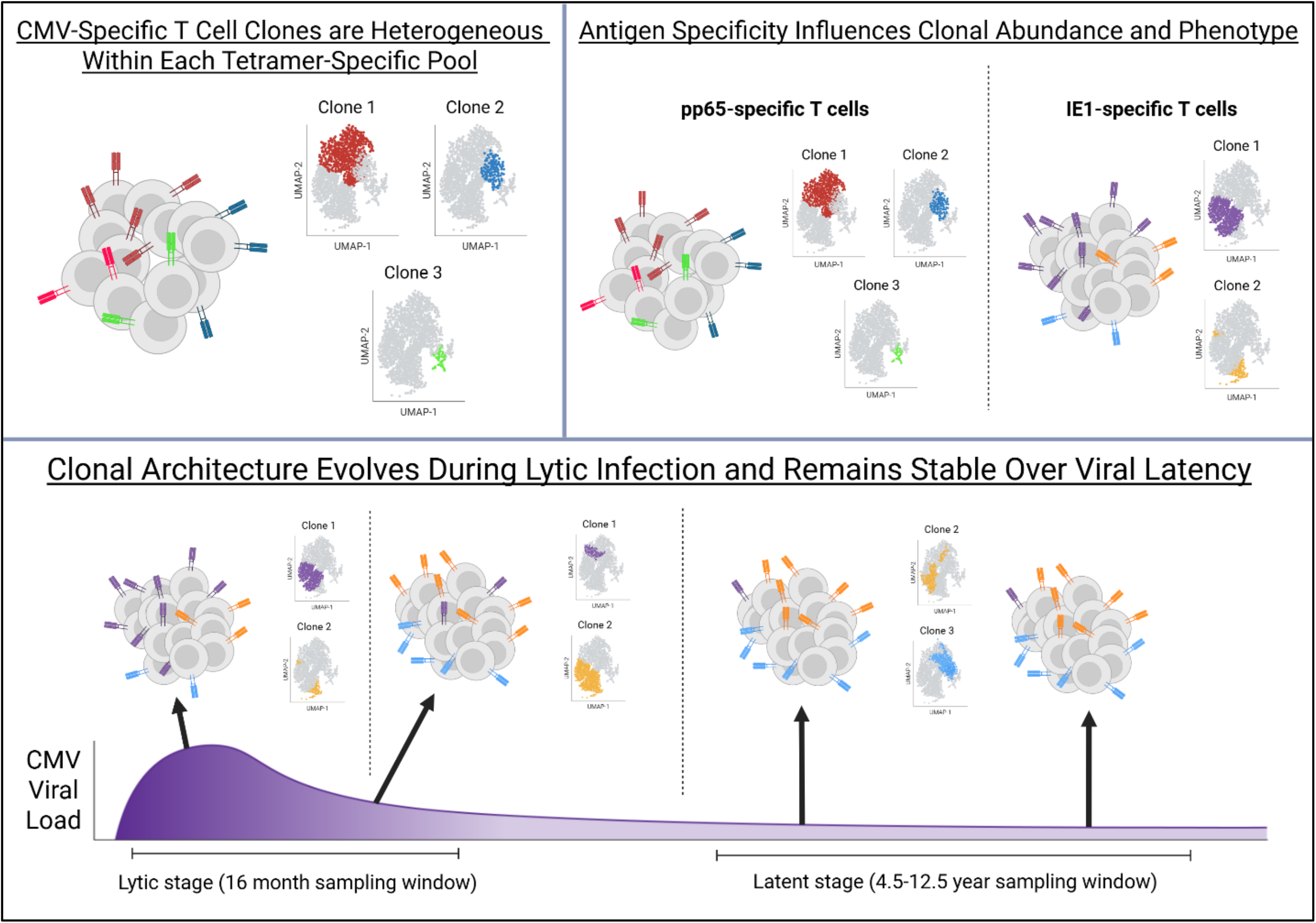

## Introduction

Human cytomegalovirus (CMV) is a betaherpesvirus and one of the most common viruses in the world, with seroprevalence exceeding 50% in adults in the United States and approaching 90% in some populations^1^. CMV is also a latent virus, and after the initial viremic infection is controlled, the virus persists indefinitely within the host, undergoing periodic, subclinical reactivation events that are largely suppressed by CD8^+^ T cell-mediated immunity^2–7^. This chronic antigen exposure drives a remarkably robust and durable T cell response in humans, with CMV-specific CD8^+^ T cells maintained at high frequencies for decades. In some elderly individuals CMV-specific T cells expand to constitute over 50% of the entire circulating CD8^+^ T cell compartment in a phenomenon known as "memory inflation"^8,9^. The magnitude and persistence of this response has caused CMV seropositivity to be consistently identified as one of the primary factors influencing immune phenotype in humans^10,11^. In particular, the aging of the immune system, or “immunosenescence”, is highly correlated with CMV infection status^9,11^, with immunosenescent individuals experiencing higher susceptibility to infectious diseases, lower vaccine responses, and higher cancer risk^9,12^.

As a result of the chronic, periodic antigenic stimulation experienced by CMV-specific T cells, these cells exhibit phenotypic and functional characteristics associated with T cell senescence, including upregulation of CD45RA, CD57, and PD-1^6,8,13,14^. Yet paradoxically, CMV-specific T cells remain functionally competent, retaining the ability to produce effector cytokines and cytolytic molecules and effectively controlling viral reactivation^6,7^. This functional resilience in the face of repeated antigenic stimulation makes CMV-specific T cells a critical model for understanding how CD8^+^ T cells maintain effector functions under conditions of chronic activation, a question with direct relevance to cancer immunotherapy, chronic viral infections, and autoimmunity.

CMV-specific T cells have also garnered significant interest for their therapeutic potential. The high frequency of these cells in some individuals’ circulation, in addition to their ability to retain functionality in the face of chronic antigen exposure, has led to attempts to re-target these cells against cancer by loading the cancer cells with immunogenic CMV peptides^15–17^. For the same reasons, CMV-specific T cells are considered a potentially attractive platform for chimeric antigen receptor (CAR) or engineered T cell receptor (TCR) cell therapies^18,19^. Additionally, CMV’s ability to drive sustained high-frequency T cell responses has led to the development of CMV-based vaccines, using a modified version of the virus expressing a vaccine target antigen in place of a naturally inflation-driving epitope^5,20^. This approach is particularly promising for HIV, where driving a strong virus-specific effector CD8^+^ T cell population would be a highly desirable vaccination outcome^21,22^. Understanding how CMV-specific T cell populations evolve over time and retain functionality in the face of chronic stimulation is therefore critical to inform strategies utilizing these cells across multiple therapeutic contexts.

Much of our current understanding of the dynamics of CMV-specific CD8^+^ T cell responses comes from longitudinal studies in mouse models of murine CMV (MCMV) infection. During acute MCMV infection CD8^+^ T cell responses are dominated by cells specific for certain viral proteins, such as M45, with higher avidity TCRs driving the most dominant clonal populations. However, during latency, the response becomes dominated by T cells specific for a different set of proteins including M38 and m139, with more low- and medium-avidity TCRs found in the dominant clones^23^. In addition, once the dominant clones are established in the latent phase of MCMV infection, these clones are generally stable for the remainder of the mouse’s life^6,23^. Mouse models also reveal that inflationary MCMV-specific T cell populations are maintained by continuous production of short-lived effector cells, the product of lymph node-resident memory populations receiving chronic antigenic stimulation^24^.

Mouse models of MCMV infection are limited by the lifespan of the animal, with studies terminating after at most 1-2 years, whereas in human CMV infection T cells may continue to evolve in the context of chronic stimulation for several decades. Human CMV-specific T cells are also very phenotypically diverse. Although the majority of these cells exhibit an effector memory (Tem) or CD45RA-re-expressing effector memory (T_EMRA_) phenotype, smaller central memory (Tcm) populations also consistently observed^25^. In a subset of individuals, some CMV-specific T cells upregulate genes typically associated with NK cells, including *KLRC2* and *TYROBP*, and are able to be activated through the NK cell activating receptor NKG2C^26^. The vast majority of human studies of human CMV-specific T cells have been cross-sectional, capturing T cell phenotypes at single timepoints during latency but providing little insight into how these phenotypes are established, or whether they remain stable over time within individuals. Studies that have tracked human CMV-specific T cells over time found that TCR clonality is preserved over up to 5 years^27,28^. However, since most studies of human CMV-specific T cells have not included single-cell TCR sequencing (scTCRseq) it remains unclear whether the phenotypic diversity in CMV-specific T cells is clonally restricted. Additionally, the majority of human studies have focused on pp65 (specifically the HLA-A*02-restricted NLVPMVATV peptide) as the main target of the T cell response, as it is one of the most immunodominant viral epitopes^29^. However, studies looking at multiple antigens, as well as the murine data on different MCMV antigens, indicates that CD8^+^ T cell responses specific for other antigens could be distinct from pp65-specific cells in terms of kinetics, clonality, and phenotype^7,23^.

To address these knowledge gaps, we used single-cell RNA sequencing (scRNAseq) with paired scTCRseq and CITEseq to profile MHC tetramer-isolated pp65- and IE1-specific CD8^+^ T cells from an acutely infected seroconverting donor and a cohort of five latently infected serial blood donors. Our analyses revealed that during the first 6 months after infection, CMV-specific T cell kinetics follow a canonical pathway of effector cell differentiation, with little change over time in the clonal architecture or transcriptional phenotype of individual clones. Between 6 months to 1.5 years post-infection, the CMV-specific T cell pool contracted, undergoing a reduction in clonal diversity and a dramatic shift in clonal transcriptional phenotype, based on antigen specificity. From our latently infected cohort, we determined that CMV-specific T cell pools were highly stable with minimal clonal turnover in most donors even over intervals exceeding a decade. These responses were phenotypically diverse, with antigen specificity once again determining both clonal architecture and transcriptional signatures. Transcriptional diversity was clonally encoded, with individual clones expressing distinct phenotypes that we classified as central memory-like, effector memory-like, NK-cell-like, and a newly described population characterized by high expression of the mitochondrial chaperone protein CMC1. We further found that relative TCR avidity was one driver of clonal abundance, phenotype, and proliferative capacity, and TCR sequence similarity drove clones to similar differentiation states across multiple individual donors. Together, our findings demonstrate that the diversity of CMV-specific T cells is clonally encoded, influenced by antigen specificity and TCR sequence, and highly stable over time. These results provide a framework for understanding how chronic viral infections shape durable CD8^+^ T cell responses in humans, which could improve our ability to harness these responses for anti-cancer or vaccination strategies.

## Results

### CMV-specific T cell clonal phenotypes are established over the first year of latent infection

To study the dynamics of human CMV-specific T cell responses during primary infection, we obtained serial PBMC samples from a donor who was infected during the two-year monitoring period of the Sound Life Project study^11^. This healthy subject seroconverted to CMV^+^ during a 4.5-month window between donated blood samples. Based on this subject’s HLA-type, we analyzed CMV-specific CD8^+^ T cell responses in samples spanning 16 months in total, using fluorescent HLA-A tetramers presenting known CMV epitopes derived from the pp65 (HLA-A*01:01/YSEHPTFTSQY) and IE1 (HLA-A*24:02/AYAQKIFKIL) proteins. These tetramers revealed populations of pp65- and IE1-specific CD8^+^ T cells which appeared in the first post-seroconversion visit at 0.27% and 3.58% of CD8^+^ cells respectively, remained relatively stable over the next 3 months, and substantially decreased in frequency during the next 9 months (Fig 1A), mirroring the clonal expansion and contraction observed in CD8^+^ T cell response to vaccination or non-latent viral infection.

**Figure 1.**
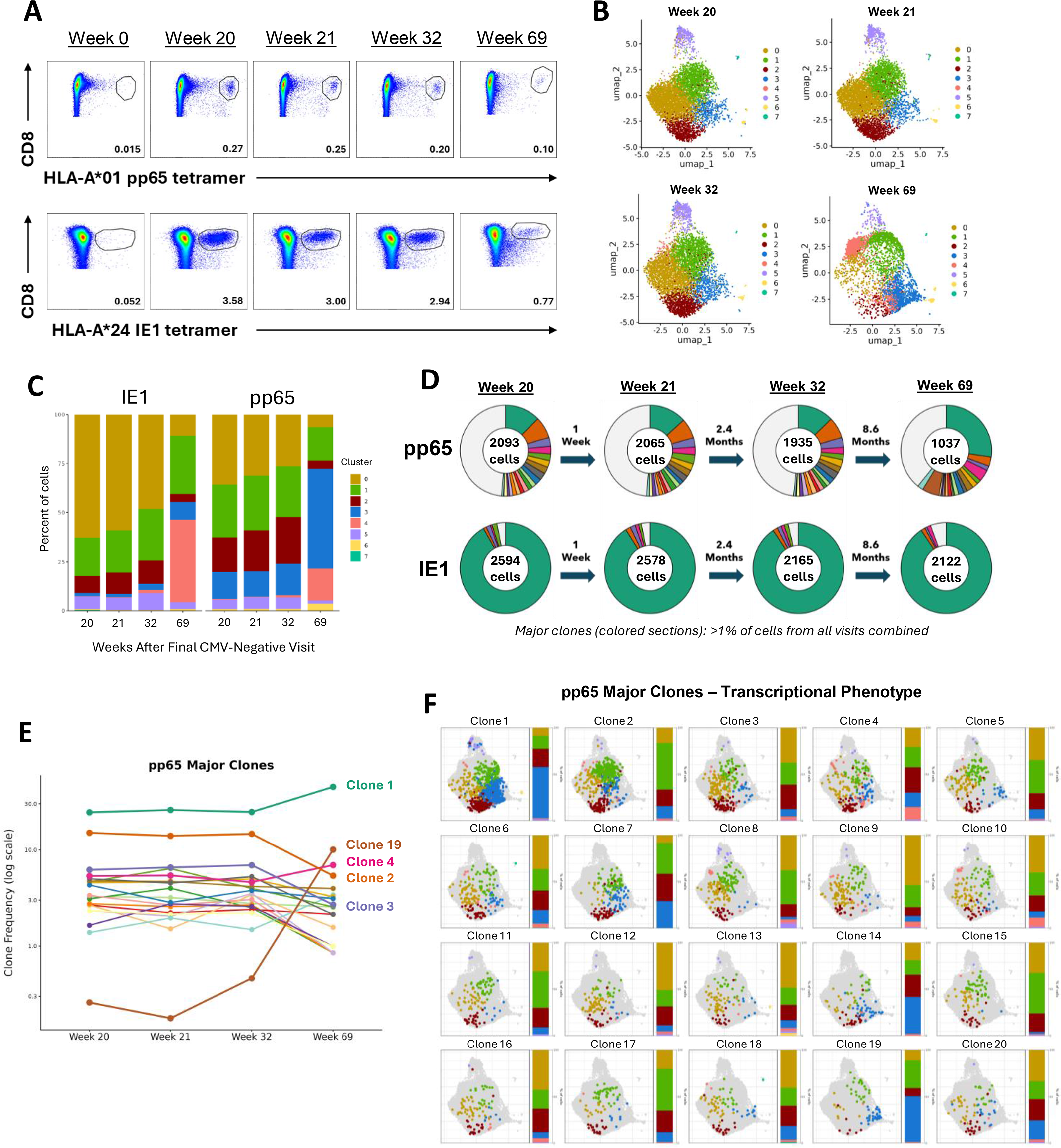
pp65- and IE1-specific T cell clonal phenotypes are established over the first year of CMV infection. **A.** Flow cytometry staining of CMV tetramers in PBMCs from the seroconverting donor across five serial blood draws. Top row, HLA-A*01 pp65 (YSEHPTFTSQY) tetramer vs CD8; bottom row, HLA-A*24 IE1 (AYAQKIFKIL) tetramer vs CD8. Numbers in each gate indicate the frequency of tetramer-binding cells as a percentage of parent population (CD8^+^ T cells). **B.** UMAP of scRNAseq data from sorted pp65- and IE1-tetramer-binding CD8^+^ T cells, split by visit. **C.** Stacked bar plots showing the percentage of cells in each Seurat cluster at each visit, split by tetramer specificity. **D.** Donut plots of clonal composition of the pp65- and IE1-specific pools at each visit. Colored wedges are major clones, defined as clonotypes comprising >1% of cells from all visits combined; the pooled grey wedge represents all remaining minor clones. **E.** Frequency of pp65-specific major clones as a percent of pp65-tetramer-binding cells, across visits, plotted on a log scale. Clones which either begin or end as one of the top 3 most dominant are highlighted. **F.** UMAP position of each pp65-specific major clone, showing data from all four post-seroconversion visits combined. Each panel shows cells belonging to that clone colored by Seurat cluster; the adjacent stacked bar graph shows the cluster composition percentages.

To assess the functional differentiation and clonal dynamics of the CD8^+^ T cell response during acute infection, we performed 10X single cell RNA sequencing (scRNAseq) with CITEseq feature barcoding and paired TCR sequencing on pp65- and IE1-specific T cells from each post-infection visit, using Uniform Manifold Approximation and Projection (UMAP) dimensionality reduction to visualize the transcriptional data (Fig 1B). Louvain clustering identified 8 distinct clusters within the CMV-specific CD8^+^ T cells, 6 of which represented major populations that together contained >95% of pp65- and IE1-specific cells. For the first three post-infection visits up to week 32, the majority of CMV-specific T cells were found in clusters 0, 1, and 2 (Fig 1B, Fig S1A). These three clusters had gene expression patterns indicative of effector T cells, including expression of the HLA-DR genes *HLA-DRB1* and *HLA-DRA* (clusters 0 and 1), cytokines and chemokines produced by activated T cells such as *TNF, CCL3*, *CCL4* and *XCL1* (cluster 2), the chemokine receptor *CX3CR1* (all three clusters), as well as surface expression of CD57 detected by CITEseq in clusters 0 and 1 (Fig S1B-C). Concurrent with their contraction, by the final visit at week 69 the cells had redistributed and the majority of cells were found in clusters 1, 3 and 4. Cluster 3 was characterized by expression of genes associated with T cell memory (*IL7R*, *TCF7*, *GZMK*) and expressed high levels of CD45RA, CXCR3, and PD-1 by CITEseq, whereas cluster 4 expressed genes associated with suppression of T cell activation (*LILRB1*, *AKT3*) along with high levels of CD57 and CD226 (Fig S1B-C). Throughout this course of early infection, pp65- and IE1-specific T cells were transcriptionally distinct, with pp65-specific cells being over-represented in cluster 2 at weeks 20-32 and cluster 3 at week 69, and IE1-specific cells over-represented in cluster 0 at weeks 20-32 and cluster 4 at week 69 (Fig 1C). This differential clustering corresponded to lower CD57 and higher CXCR3 in the pp65-specific cells, particularly at week 69 (Fig S1D). Analysis of TCR sequences in pp65- and IE1-specific T cells revealed a dramatic difference in TCR clonality between the two populations. We assigned all cells into clonotypes based on paired TCRα and TCRβ CDR3 nucleotide sequences. Although the size of the IE1-specific T cell pool was >10x larger than that of the pp65-specific pool based on percentage of total CD8^+^ T cells at the peak of the response (Fig 1A), we observed that over 90% of the IE1-specific cells at all timepoints examined belonged to a single TCR clone (Fig 1D). In contrast, pp65-specific cells were much more clonally diverse, with the most dominant pp65-specific clone making up around 25% of that pool and 19 sub-dominant major clones (consisting of at least 1% of all sampled tetramer-binding cells) identified. These pp65-specific major clones made up 50-60% of the response at each timepoint, with the remaining 40-50% made up of ∼900 clones each comprising <1% of tetramer-binding cells. Across the first three visits (spanning 2.5 months), the relative abundance of pp65-specific clones was highly stable (Fig 1E). However, although all 20 major pp65-specific clones were still present at week 69, each followed its own contraction trajectory resulting in a shift in the clonal hierarchy, with some pp65-specific clones that were less abundant during the initial visits (clones 4 and 19) growing to represent over 5% of the response, and others that were more dominant (clones 2 and 3) shrinking as a fraction of the overall pool (Fig 1E, Fig S1E-F). This shift corresponded to a significant decrease in clonal repertoire heterogeneity of the pp65-specific response as measured by Gini-Simpson index, although it was still much more diverse than the IE1-specific response (Fig S1G).

We also observed a high level of transcriptional heterogeneity amongst pp65-specific clones, with individual clones showing unique patterns of gene expression in UMAP space and distribution among the effector and memory clusters (Fig 1F). Although there were far fewer IE1-specific clones, the dominant clone also showed a unique expression pattern compared to the sub-dominant clones (Fig S2A). This transcriptional heterogeneity corresponded to heterogeneity of CITEseq marker expression between clones as well, especially in CD57, CXCR3 and CD226 (Fig S2B). We then considered the differentiation trajectory of each clone across the four visits, by calculating the Jensen-Shannon divergence of each clone’s distribution amongst the 8 clusters compared to its distribution at week 20 (Fig S2C). This showed that clonal phenotypes were mostly stable for weeks 20-32 but had a dramatic phenotypic remodeling by week 69. This remodeling generally followed the path of moving from a cluster 0/1/2-like phenotype to a cluster 1/3/4-like phenotype, but individual clones were highly variable in their differentiation trajectories among clusters during contraction (Fig S2D). Overall, these analyses revealed that transcriptional and cellular differentiation following primary infection is associated with antigen specificity while also varying substantially among individual clonotypes. Clonal phenotypes were established early after infection and subsequently followed distinct trajectories during contraction and memory formation.

### pp65- and IE1-specific T cells are phenotypically distinct in latently infected donors

To more generally study the phenotypic, transcriptional, and clonal stability of CMV-specific CD8^+^ T cell populations during viral latency, we identified a cohort of individuals in the BRI Immune-Mediated Disease Repository who had donated blood for research purposes multiple times over the course of several years. This criteria identified 150 serial donors, including healthy control donors and donors with type 1 diabetes who were not on immune-modifying therapies. These individuals all expressed the common HLA allele HLA-A*02, which presents dominant peptide antigens from both pp65 (NLVPMVATV) and IE1 (VLEETSVML), allowing us to use MHC-tetramers to identify CMV-reactive cells specific for the same antigenic proteins as in the acutely infected subject. Of the 150 serial donors, we found 18 whose PBMCs had detectable T cell populations binding both the pp65 and IE1 tetramers, indicating latent CMV infection. The frequencies of these populations ranged from 0.001% to 11% of all CD8^+^ T cells, with neither epitope eliciting a consistently larger response (Fig 2A). This cohort ranged in age from 26-69 and we did not observe a significant correlation between age and the size of the CMV-specific T cell populations (Fig S3A).

**Figure 2.**
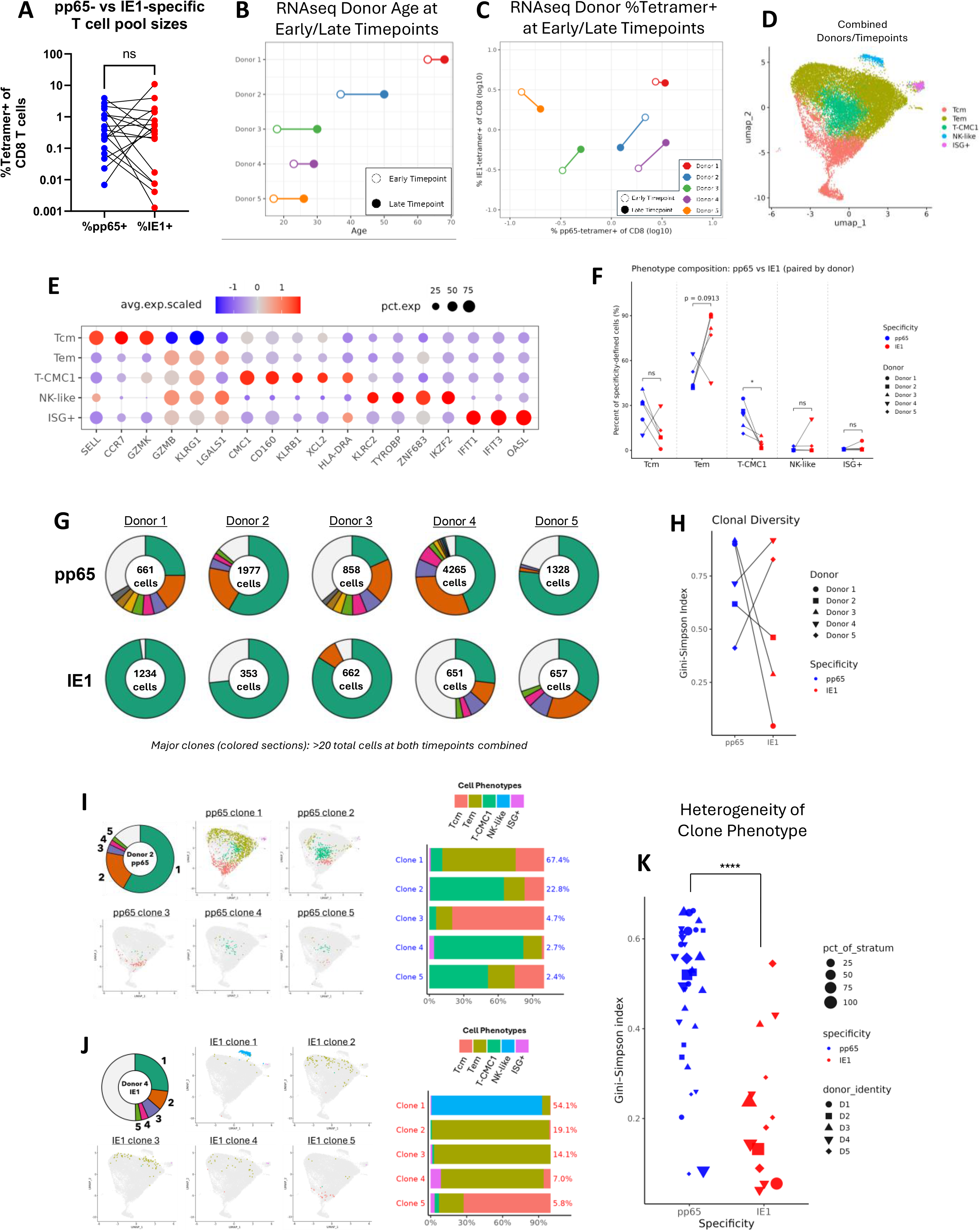
Specificity-dependent differences and clonal heterogeneity within latently infected individuals. **A.** Frequency of pp65- and IE1-tetramer-binding cells as a percentage of total CD8^+^ T cells in the latently infected screening cohort, with paired samples from the same donor connected. **B.** Age of each scRNAseq donor at the early and late timepoint analyzed. **C.** Size of the pp65-specific (x- axis) and IE1-specific (y-axis) T cell pools, as a percentage of CD8^+^ T cells, at the early and late timepoint for each scRNAseq. **D.** UMAP of integrated scRNAseq data from all tetramer-binding CMV-specific CD8^+^ T cells, pooled across all five donors and both timepoints, colored by transcriptional phenotype. **E.** Dot plot of selected marker gene expression, z-score scaled across the five phenotypes. Dot size indicates the percentage of cells in the phenotype expressing the gene. **F.** Percentage of pp65-vs IE1-specific cells falling into each phenotype, paired within donor. **G.** Donut plots of clonal composition of the pp65- and IE1-specific pools for each of the five donors. Colored wedges are major clones, defined as clonotypes with at least 20 cells from both timepoints combined; the pooled grey wedge represents all remaining minor clones. **H.** Gini-Simpson index of clonal diversity for the pp65- and IE1-specific pools of each donor, paired within donor. **I.** Transcriptional phenotype of the top five pp65-specific clones from donor 2. Left, donut plot of the pp65-specific pool with the top five clones numbered. Center, UMAP position of each clone, colored by phenotype. Right, stacked bar of phenotype composition for each clone; the value to the right of each bar gives that clone’s percentage of the pp65-specific pool. **J.** As in (I), for the top five IE1-specific clones from donor 4. **K.** Gini-Simpson index of diversity of phenotypes within each clone (phenotypic heterogeneity) for each major clone, grouped by antigen specificity. Point shape indicates donor and point size indicates the clone’s percentage of its antigen-specific pool. *p < 0.05, **p < 0.01, ***p < 0.001, ****p < 0.0001.

A flow cytometry-based screening assay of these donors revealed that IE1-specific T cells had higher surface levels of CD57 than did pp65-specific T cells, mirroring our CITEseq results from the seroconverting donor (Fig S3B). Within IE1-specific T cells, CD57 expression was positively correlated with the size of the epitope-specific T cell pool (Fig S3C). IE1-specific T cells also expressed significantly lower levels of CD27, indicating a more effector-like state (Fig S3D). This corresponded to significantly lower frequencies of CD27^+^CD45RA^+^ and CD27^+^CD45RA^-^cells, as well as significantly higher frequencies of CD27^-^CD45RA^-^cells and a non-significant trend toward higher levels of CD27^-^CD45RA^+^ T_EMRA_ cells (Fig S3E-F). After stimulation with cognate antigen, pp65-specific cells upregulated significantly higher levels of the activation-induced markers CD25 and CD137 compared to IE1-specific cells (Fig S3G-H).

#### scRNAseq of CMV-specific CD8^+^ T cells in longitudinal PBMC samples from five donors

To determine how pp65- and IE1-specific T cell responses evolve during the latent phase of CMV infection, we selected five subjects from our cohort based on high frequency of detectable pp65-and IE1-specific T cells and the availability of frozen PBMC samples spanning more than four years. We performed scRNAseq with TCRseq and CITEseq on sorted populations of pp65-specific, IE1-specific, and non-tetramer-binding bulk CD8^+^ T cells, from two selected blood draws for each donor (Fig S4A, Table S1). To ensure these donors had not been recently infected, CMV tetramer-binding cells and/or CMV IgG were confirmed to be present in PBMCs/serum from at least one year prior to the first selected time point in donors 1, 3, and 5. For donor 4, no earlier samples were available to test, however CMV serology from the first selected sample showed that this donor was CMV IgG+ and IgM-, indicating that their CMV infection was likely already into the latent stage by this time^30^. Donor 2 was also tested to be CMV IgG^+^ at the first selected sample (Table S1). The five selected donors ranged in age from 17 to 63 at the time of the first blood draw analyzed, with 4.5 to 12.5 years between the two tested samples (Fig 2B). The pp65-specific T cell populations represented 0.13% to 3.45% of total CD8^+^ T cells, and the IE1-specific populations represented 0.31% to 3.98% of CD8^+^ T cells, with no consistent increase or decrease in the frequencies of either specificity over time across donors (Fig 2C).

UMAP dimensionality reduction on integrated transcriptional data from all donors and timepoints showed differential RNAseq expression patterns between tetramer-negative cells, pp65-tetramer-binding cells, and IE1-tetramer-binding cells (Fig S4B). Tetramer-negative cells were found throughout the UMAP space, but selectively enriched relative to pp65- and IE1-specific cells in regions associated with naïve (*CCR7*^+^) and MAIT (*ZBTB16*^+^) cell markers, while tetramer-binding cells were concentrated into the *KLRG1^+^* region (Fig S4C). IE1-specific cells in particular were concentrated in the *GZMB^+^*region and also the *TYROBP*^+^ NK-like cell region, whereas pp65-specific cells were found across the *KLRG1*^+^ region expressing either *GZMB* or *GZMK* (Fig S4B-C).

### Transcriptional phenotype differs based on antigen specificity

To further assess the heterogeneity within each donor based on specificity for either pp65 or IE1, we subsetted our scRNAseq data to include only tetramer-binding CMV-specific cells. Unbiased Louvain clustering in Seurat showed nine distinct clusters of cells within this dataset (Fig S5A). Based on differentially-expressed genes (DEGs), we assigned these clusters into phenotypic categories, which included Tcm (clusters 3 and 4), Tem (clusters 0, 2, 7 and 8), NK-like (cluster 5), ISG^+^ (cluster 6), as well as a newly described T cell phenotype T-CMC1 (cluster 1) (Fig S5B-C)^31,32^. These T-CMC1 cells lie between the Tcm and Tem populations in UMAP space (Fig 2D) and were marked by high expression of the gene *CMC1*, as well as certain NK-cell-associated genes such as *CD160* and *KLRB1* (Fig 2E). T-CMC1 cells also showed expression of the activation markers *HLA-DRA* and *HLA-DRB5* (Fig S5B). Our CITEseq data revealed that T-CMC1 cells also had the highest expression of both CD45RA and PD-1 (Fig S5D). Compared with IE1-specific T cells, pp65-specific cells had a significantly higher percentage of the T-CMC1 phenotype, as well as a trend towards increased Tcm phenotype and decreased Tem phenotype in 4 of the 5 analyzed donors (Fig 2F). The NK-like and ISG^+^ phenotypes were mostly found in IE1-specific cells, and the majority of these populations were found in a single donor in both cawses (Fig 2F).

This observed trend of pp65-specific cells being Tcm-biased, and IE1-specific cells being Tem-biased, mirrored the specificity-dependent differences we observed in our seroconverting donor. A direct comparison of DEGs between pp65- and IE1-specific cells in the two datasets showed that 88% of genes which were up- or down-regulated in pp65 vs IE1-specific T cells in the latently infected cohort were also up- or down-regulated in cells of the same specificity in the seroconverting donor (Fig S5E). Even when only considering that donor’s early timepoints (the first 3 visits from week 20-32), we still observed an 85% DEG overlap, including Tcm-vs-Tem defining genes such as *IL7R*, *TCF7*, and *GZMB* (Fig S5E).

To place our transcriptional findings in the broader context of chronic T cell stimulation, we compared our data with published studies of T cells in cancer using a published transcriptomic atlas of CD8^+^ T cells from PBMCs, healthy tissue, and cancer biopsies^33^. This atlas defined 17 clusters of CD8^+^ T cells ranging from naïve to terminally exhausted (Fig S6A). By generating gene sets from the top DEGs of each of our 5 phenotypes and applying them to the published dataset, we observed that our NK-like, Tem, Tcm, and ISG+ phenotypes broadly recapitulated those defined in the atlas, overlapping with the KIR^+^, Temra, *IL7R*^+^ Tm, and *IFIT*^+^ ISG clusters respectively (Fig S6B). Interestingly, the T-CMC1 phenotype had the highest overlap with 2 of the exhausted T cell (Tex) clusters, as well as the NK-like cluster (Fig S6B). This finding aligns with the limited prior studies that identified CMC1 as a marker of dysfunctional T cells in the context of tumor immunology^31,32^.

### Transcriptional heterogeneity of CMV-specific T cells is clonally encoded

As in our analysis of the acutely infected subject, we used TCRseq data to assign pp65- and IE1-specific T cells into dominant and sub-dominant clonotypes within each donor, defining major clones as antigen-specific cells with the same TCR V, D, and J gene usage and identical TCRα/TCRβ CDR3 nucleotide sequences, with at least 20 cells detected across both timepoints (ensuring we had enough power to define the clones’ phenotypes). We observed that in four out of the five donors, the response against one antigen was oligoclonal while the response against the other was >70% dominated by a single clone (Fig 2G). Of these four, three had near-monoclonality in the IE1-specific pool, similar to what we observed in the acutely infected donor (Fig 1D). However, across all five latently infected donors we observed no significant difference in the clonality of IE1- and pp65-specific responses (Fig 2H).

As we found during acute infection, transcriptional heterogeneity within CMV-specific T cells in our latent cohort was clonally encoded (Fig S7). For example, the top five pp65-specific clones from donor 2 had a clone with a majority Tem phenotype (clone 1), a clone with a majority Tcm phenotype (clone 3), and three clones with a majority T-CMC1 phenotype (clones 2, 4, and 5) (Fig 2I). Similarly, the top five IE1-specific clones from donor 4 had a majority-NK-like clone, three majority-Tem clones, and a majority-Tcm clone (Fig 2J). Overall, pp65-specific clones had a significantly higher phenotypic diversity than did IE1-specific clones, as defined by Gini-Simpson index of the spread of each clone amongst the five phenotypes (Fig 2K). We also observed that the relative size of each clone within its antigen-specific pool did not correlate with any specific transcriptional phenotype, or with the clone’s phenotypic diversity (Fig S7, Fig 2K).

### Clonal Dominance and Transcriptional Phenotype are Stable Over Time

In 4 of 5 donors, clonal dominance hierarchies among both pp65- and IE1-specific populations were stable across the two timepoints analyzed, ranging from 4.5-12.5 years apart (Fig 3A-B). However, for donor 3, although we did detect the same major clones appearing at both timepoints, there was a dramatic shift in dominance among the top clones for both specificities. Nevertheless, across the 48 major clones of CMV-specific T cells from all donors, the frequency of each clone at the early timepoint was significantly correlated with its frequency at the late timepoint (Fig 3C), with clones from donor 3 being the primary outliers to this correlation. Since donor 3 had been infected with CMV for at least 1.5 years at the point of their first donation, this deviation cannot likely be explained by that donor being sampled during the lytic-to-latent transition stage.

**Figure 3.**
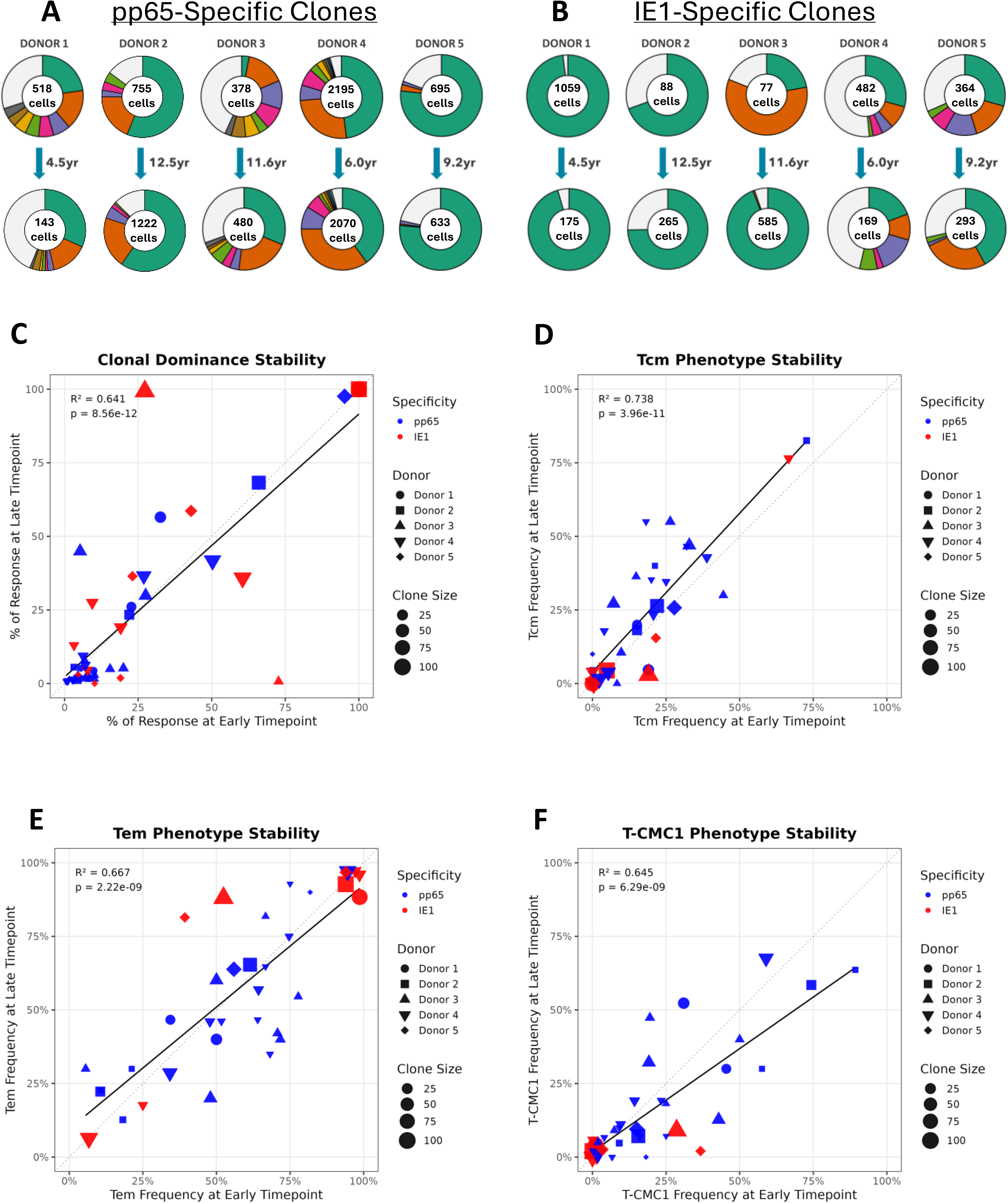
Clonal dominance and transcriptional phenotype are stable over time. **A.** Donut plots of pp65-specific clonal composition at the early (top) and late (bottom) timepoint for each of the five donors. Colored wedges are major clones; wedge colors are matched between the early and late plot within each donor. **B.** As in (A), for IE1-specific clones. **C.** Correlation between each major clone’s percentage of its antigen-specific pool at the early vs the late timepoint. Point color indicates specificity, shape indicates donor, and size indicates the clone’s percent of antigen-specific response across both timepoints. Solid line with R^2^ and p-value represent linear regression. **D-F.** Correlation between the percentage of cells within each clone assigned to a given phenotype at the early vs the late timepoint, for Tcm (D), Tem (E), and T-CMC1 (F). Points and lines as in (C). Only clones with at least 20 cells across both timepoints are included.

We also calculated the percentage of cells falling into each functional phenotype (Tcm, Tem, NK-like, ISG^+^, or T-CMC1) for all clones with at least 20 cells from both timepoints (Fig S8A). We found that clonal phenotypes were highly stable for most clones over the 4.5-12.5 year gaps between samples (Fig 3D-F). For the NK-like and ISG^+^ phenotypes, there were almost no clones with an appreciable frequency in either phenotype, with the only exception being a dominant IE1-specific clone in donor 4, which was almost entirely NK-like at both timepoints (Fig S8B-C). Overall, we observed a remarkable level of stability in both relative dominance and transcriptional phenotype of most clones during long periods of latent infection and potential attempted viral reactivation, even amongst highly heterogeneous and oligoclonal populations.

### Relative TCR Affinity Influences Clonal Proliferative Capacity and Phenotype

To determine if each clone’s abundance and distinct phenotypes may be due to their relative avidity for antigen, we stained CD8^+^ T cells with a limiting-dilution of pp65 or IE1 tetramer and sorted high-vs low-MFI tetramer binding cells (Fig 4A). We then performed bulk TCR seq on each population to find the frequencies at which each clone’s TCR was recovered from both sort gates, as well as a control total tetramer^+^ sort (Fig 4B). After normalizing the percent abundance of each TCR sequence in each sorted population using the centered-log ratio (CLR), we calculated a relative TCR avidity score for each clone based on the CLR-normalized abundance of that clone’s TCR in the high-MFI vs low-MFI tetramer populations (Fig 4C). Due to limiting sample numbers, we tested pp65-specific clones from donors 1-4 and IE1-specific clones from donors 4 and 5, based on these being the most polyclonal responses against each antigen. To confirm that the differential enrichment we observed between high-MFI and low-MFI sorted fractions reflected genuine differences in TCR avidity rather than stochastic variation in clonal sampling, we repeated the sorting and sequencing for another PBMC sample from one of the tested draws, and we found that clonal enrichment scores were highly consistent across technical replicates (Fig S9A).

**Figure 4.**
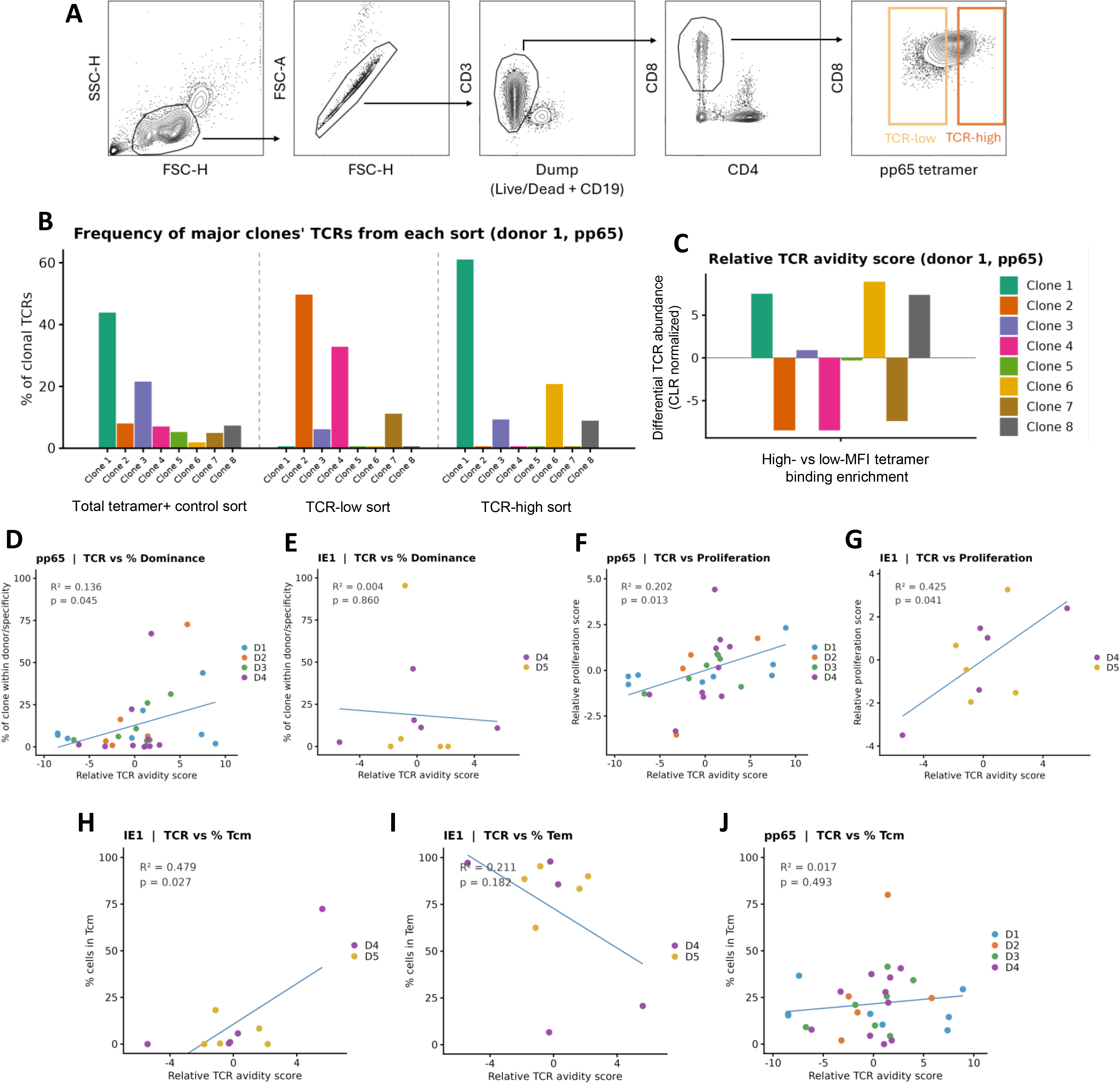
Relative TCR affinity influences clonal proliferative capacity and transcriptional phenotype. **A.** Example gating strategy for sorted populations: singlets, CD3^+^, CD8^+^, pp65 tetramer-high and low MFI binding. **B.** Frequency at which each clonal TCR from donor 1’s pp65-specific major clones appeared in the total tetramer+ control, TCR-low, and TCR-high sort gates. Data shown as the percentage of UMIs sequenced corresponding to each clone, relative to the total number of UMIs from all major clones. **C.** Relative TCR avidity score for each of donor 1’s pp65-specific major clones, calculated as the CLR-normalized abundance in the TCR-high sort minus CLR-normalized abundance in the TCR-low sort. **D.** Relative TCR avidity score vs each pp65 clone’s percentage of the pp65-specific pool within its donor, colored by donor, with linear regression. **E.** Relative TCR avidity score vs each IE1 clone’s percentage of the IE1-specific pool within its donor. **F.** Relative TCR avidity score vs relative proliferation score for pp65-specific major clones. **G.** Relative TCR avidity score vs relative proliferation score for IE1-specific major clones. **H.** Relative TCR avidity score vs percentage of cells with the Tcm phenotype, IE1-specific clones. **I.** Relative TCR avidity score vs percentage of cells with the Tem phenotype, IE1-specific clones. **J.** Relative TCR avidity score vs percentage of cells with the Tcm phenotype, pp65-specific clones.

Relative TCR avidity had a significant positive correlation with the clonal frequency of pp65-specific clones, but not IE1-specific clones (Fig 4D-E). To explore this further, we also wanted to determine if individual clones had differing proliferative capacity corresponding to their TCR avidity or clonal frequency. To that end, we sorted CD8^+^ T cells after a 5-day proliferation assay and used bulk TCR seq, calculating a relative proliferation score based on each clonal TCR’s CLR-normalized abundance in the proliferated sample compared to control sample (Fig S9B-C). We found that clonal proliferation score had a significant positive correlation with relative TCR avidity score for both pp65-and IE1-specific clones (Fig 4F-G). These results indicate that higher avidity clones within each response have a slight proliferative edge against their peers, which may correspond to increased clonal dominance for pp65-specific cells.

When comparing the scRNAseq profile of clones to their relative TCR avidity and proliferation scores, we observed a significant positive correlation between relative TCR avidity of IE1-specific clones and percent of cells in that clone with the Tcm phenotype (Fig 4H). We also observed a trend towards a negative correlation between relative TCR avidity score and percent of cells with the Tem phenotype for IE1-specific clones (Fig 4I). For pp65-specific clones, there was no correlation between relative TCR avidity score and either Tcm or Tem phenotype (Fig 4J, Fig S9D), but we did observe a trend towards negative correlation between relative TCR avidity score and T-CMC1 phenotype (Fig S9E-F). These findings support the idea that the strength of the interaction between TCR and peptide-MHC impact each clone’s relative abundance, differentiation trajectory, and phenotype^34^.

### Convergent Clonal TCR Sequences Appear Across Donors and Correlate with Phenotype

T cell differentiation status is partially dependent on cell-extrinsic factors, such as antigen-abundance and the presence or absence of specific co-stimulatory and cytokine cues. However, the correlations we observed with relative clonal TCR avidity indicate that the biochemical features of each clone’s specific TCR may also contribute to these dynamics. To further test this hypothesis, we used TCRdist to find groups of TCRs within our dataset that utilized the same V, D, and J genes, and had identical or nearly identical amino acid sequences in the CDR3 regions of the α and β TCR chains^35^. Of our 48 major clones, 23 were found in 5 clusters of pp65-specific TCRs and 2 clusters of IE1-specific TCRs (Fig 5A). Within the pp65-specific TCRs, the exact same TRA and TRB CDR3 amino acid sequences were found in clones from multiples donors for 4 out of 5 clusters, and TCRs within each cluster were either aa-sequence-identical or differing by only a single amino acid in either the TRA or TRB (Fig 5B, Fig S10A-D). Within IE1-specific TCRs, sequence sharing was less common (potentially reflecting fewer clones identified), although some converging sequences in both TCR chains across donors were found (Fig 5C, Fig S10E).

**Figure 5.**
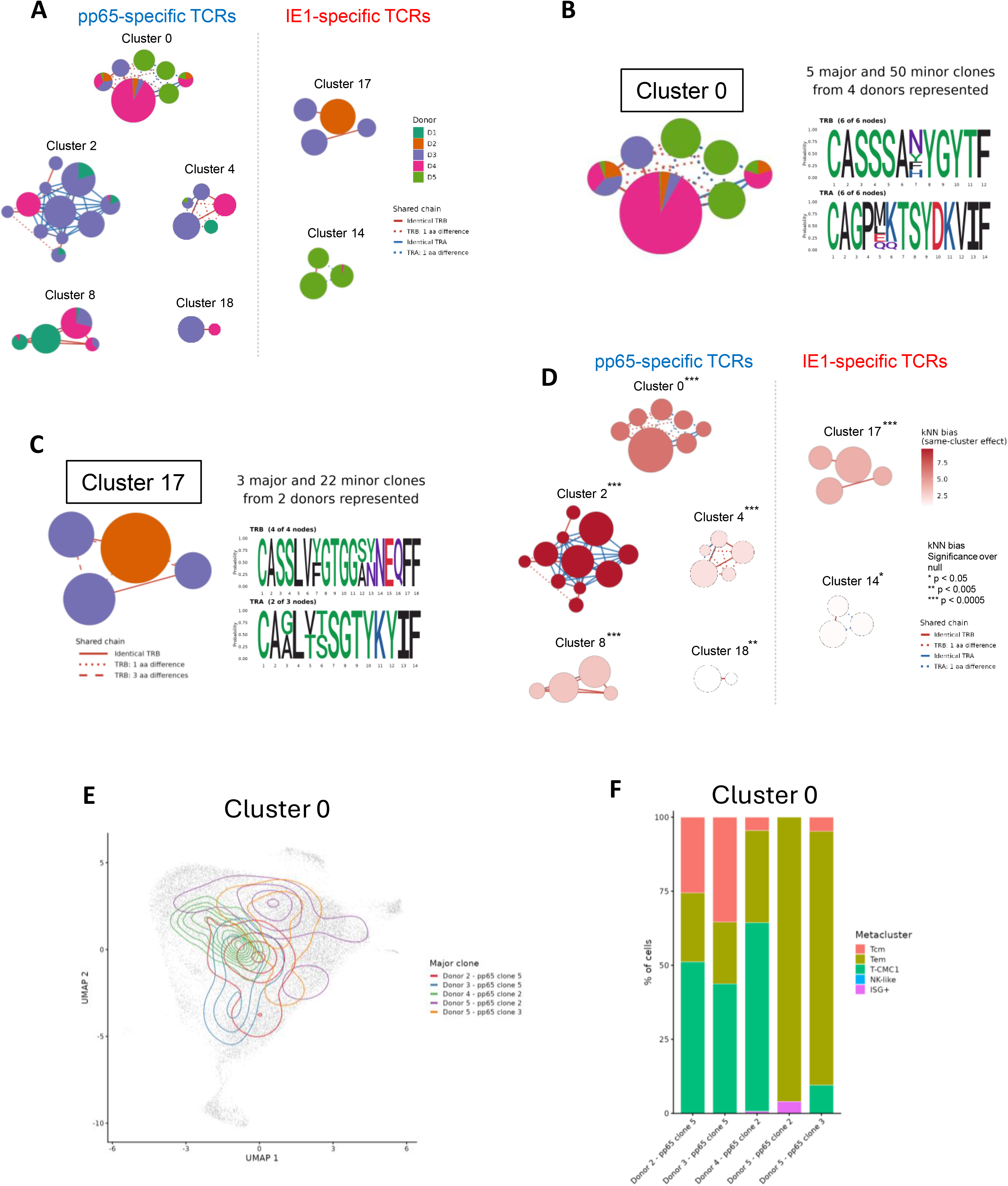
Convergent clonal TCR sequences appear across donors with shared clonal phenotypes. **A.** TCRdist-defined clusters of convergent TCRs among major CMV-specific clones. Five pp65-specific clusters (0, 2, 4, 8, 18) and two IE1-specific clusters (17, 14) are shown. Each node is a unique TRB/TRA pair; node color indicates the donor of origin (D1–D5) and pie segments indicate nodes shared between donors. Nodes are sized based on number of cells with the corresponding TCR sequence. Edges indicate the relationship between connected TCRs: solid red, identical TRB; dotted red, TRB differing by one amino acid; solid blue, identical TRA; dotted blue, TRA differing by one amino acid. **B.** Cluster 0 (pp65-specific) shown with a sequence logo of the TRB (top) and TRA (bottom) CDR3 amino acid sequences from each node. Annotation above the logo gives the number of major and minor clones and the number of donors contributing. **C.** As in (B), for Cluster 17 (IE1-specific). Dashed red line edges show TRB sequences differing by 3 amino acids. **D.** The same TCR cluster layouts as in (A), with nodes colored by k-nearest-neighbors (kNN) bias (the degree to which a clone’s neighbors in transcriptional space are enriched for cells of the same TCR cluster, relative to all other cells of the same antigen specificity). Asterisks show significance of kNN bias over the null hypothesis. **E.** UMAP density contours for each of the five major clones in TCR cluster 0. **F.** Phenotype composition of each of the five major clones in TCR cluster 0. *p < 0.05, **p < 0.005, ***p < 0.0005.

To determine if clones within a TCR cluster were phenotypically similar, we used a k-nearest neighbors (kNN) bias metric. This tested whether each clone’s neighbors in transcriptional space were more likely to be cells from the same TCR cluster compared to all other cells with the same antigen specificity. Our 7 TCR clusters were all significantly biased towards neighbors from the same cluster, with pp65-specific clusters showing a higher average kNN bias (Fig 5D). For highly biased clusters, the phenotypic similarity of clones within the cluster could also be visualized by overlapping expression in UMAP space (Fig 5E, Fig S11). For example, the clones from donors 3 and 4 with the highest percentage of T-CMC1, as well as a donor 2 clone with over 50% T-CMC1, were all found in TCR cluster 0 (Fig 5F), whereas TCR cluster 2 contained clones from donors 1, 3, and 4, all of which showed a relatively heterogeneous phenotype, with an even split between Tcm, Tem, and T-CMC1 (Fig S11A). This analysis revealed that clones from across our donor cohort converged not only in TCR sequence but also in transcriptional phenotype. That unrelated donors arrived at the same pairing of receptor and phenotype further supports the conclusion that the diversity of the CMV-specific T cell response is organized at the level of each individual clone, and indicates that this organization is imposed in part by the TCR itself.

## Discussion

The CD8^+^ T cell response to human CMV is remarkable for both its magnitude and its functional persistence in the face of chronic antigen stimulation, but the mechanisms that generate and maintain it have been difficult to resolve since most human studies to date have been cross-sectional, restricted to a single immunodominant antigen, and did not include TCR sequencing to study individual clonal states^6,7^. Here, by combining longitudinal sampling with paired scRNAseq, scTCRseq and CITEseq, we followed individual pp65- and IE1-specific clones during primary infection and through more than a decade of established latency. Three key findings emerged: First, pp65- and IE1-specific T cells show a significant phenotypic and transcriptional divergence that is established during the first 18 months of infection and preserved during latency, even across the two separate HLA restrictions and epitopes tested; second, within a single antigen specificity in a single donor, individual clones occupy distinct and heterogeneous differentiation states, and these clonal states are highly stable, remaining essentially unchanged over intervals of 4.5-12.5 years; and third, each clone’s phenotype is predicted in part by the sequence and relative avidity of its TCR, with related TCRs from within and between donors converging to similar transcriptional states.

Our analysis of a donor sampled through seroconversion and acute infection allowed us to demonstrate that both the transcriptional divergence of pp65- and IE1-specific T cells, as well as CMV-specific clonal heterogeneity, are both established early after infection. Across the first three visits, spanning 2.5 months and falling within at most 4.5 months of the infection event, both the pp65- and IE1-specific pools remained clonally and transcriptionally static. Between ∼6 months and 1.5 years post-infection, these populations underwent substantial contraction during which pp65- and IE1-specific pools shifted from an effector state towards two distinct memory-like states. This relatively protracted course aligns with the kinetics of CMV lytic infection clearance and establishment of latency, which can take several months^36–38^. Because nearly 12 months separated visits 3 and 4, denser sampling of additional seroconverting donors would be required to further define how clonal selection and memory programming occur during establishment of viral latency.

pp65- and IE1-specific cells differed in clonal architecture, transcriptional phenotype, surface phenotype and functional responsiveness, and they did so similarly in the acutely infected donor and in our latently infected cohort. IE1-specific cells were consistently the more effector differentiated, with higher CD57, lower CD27, higher expression of *GZMB*, *PRF1*, *GNLY* and *ZNF683*, and enrichment in the Tem and NK-like states, whereas pp65-specific cells retained *GZMK*, *TCF7* and *CD27* expression, were enriched in the Tcm and T-CMC1 states, and upregulated CD25 and CD137 more strongly following stimulation with cognate peptide. Because we observed these differences across the two different HLAs of the seroconverting vs latently infected donors (HLA-A*01/HLA-A*24 vs. HLA-A*02), it is unlikely to be a specific property of the peptide-MHC complex itself, and more likely reflects intrinsic properties of these viral proteins that influence how these antigens are encountered by CD8^+^ T cells during acute infection and viral latency. IE1 is an immediate-early gene product expressed at the beginning of the CMV infectious cycle, potentially including abortive events that never proceed to late gene expression^39,40^, whereas pp65 is not expressed until the late phase of infection, but is highly expressed, comprising ∼15% of the viral proteome^29^. These antigens also differ in that as a transcription factor IE1 is expressed in infected cells’ nuclei but is not found in the actual virion, whereas the viral tegument protein pp65 is a major structural component of CMV viral particles. The differences we observe in pp65- vs IE1-specific T cells likely reflect a combination of several factors that stem from these differences, including antigen abundance, differential APC presentation, tissue localization, and timing or frequency of antigen stimulation. Because these phenotypic differences between pp65- and IE1-specific T cells were already apparent in the early stages of primary infection, they are unlikely to arise solely from progressive differentiation driven by repeated or prolonged antigen exposure during viral latency, although such exposure may reinforce or amplify these distinct phenotypic states. This pattern is reminiscent of the distinction between epitopes described in MCMV, in which responses to certain gene products such as IE3 (homologous to the HCMV IE2) adopt a terminally differentiated effector-memory phenotype, while responses to other epitopes such as M45 (a viral ribonucleotide reductase homologous to HCMV UL45) contract into a conventional memory pool^23,41^. Antigen-driven phenotypic divergence is also not unique to CMV. Recent work in EBV found that CD8^+^ T cells specific for different viral epitopes adopt distinct differentiation states and follow distinct trajectories with advancing age^42^, suggesting that antigen-intrinsic specification of T cell fate may be a general feature of latent herpesvirus infection.

The newly described T-CMC1 population has not previously been studied in the context of virus-specific human CD8^+^ T cell responses. These cells occupy an intermediate position between Tcm and Tem in transcriptional space, co-express *CMC1* with *CD160* and the activation markers *HLA-DRA* and *HLA-DRB5*, and by CITEseq carry the highest surface levels of both CD45RA and PD-1 in our dataset. The T-CMC1 signature mapped most strongly onto two of the exhausted T cell clusters in a pan-cancer CD8^+^ T cell atlas, consistent with reports identifying *CMC1* expression as a marker of terminal differentiation and dysfunction in tumor-infiltrating T cells^31,32^. However, the full combination of features we observed is difficult to reconcile with exhaustion as conventionally defined. T-CMC1 cells were most abundant within the pp65-specific compartment, which was the more functionally responsive of the two specificities by activation-induced marker expression; they are not enriched within higher-avidity clones; and they persist at stable frequencies within individual clones for over a decade. We therefore favor the interpretation that T-CMC1 represents a distinct and durable adaptation to intermittent antigen exposure that shares a transcriptional module with tumor-associated exhaustion without sharing its functional collapse. This would make T-CMC1 cells a useful natural comparator for efforts to engineer T cell products that tolerate chronic stimulation. The T-CMC1 state may also be relevant in the context of autoimmunity, where sustained functionality in the face of repeated exposure to self-antigen could drive disease development or exacerbation. Directly defining the functional, metabolic, and epigenetic properties of T-CMC1 cells, and establishing the role of these cells in other disease contexts, is a priority for future work.

One striking observation from our data was the emergence of clonally-restricted populations of cells responding to one of the two viral antigens. Near-monoclonal IE1-specific responses occurred in three of five latently infected donors, with a near-monoclonal pp65-specific response in one other donor. Similarly, in the seroconverting donor, the IE1-specific pool outnumbered the pp65-specific pool by more than tenfold at the peak of the response, yet more than 90% of IE1-specific cells belonged to a single clonotype at every timepoint examined. Thus, clonal restriction appears to be established early, during the initial lytic infection or the transition from initial infection to latency, rather than accrued gradually by competition for limiting antigen during latency. Whether this reflects a restricted naive precursor pool, strong intraclonal competition, or the early recruitment of a single unusually fit clone cannot be resolved from our data.

The strongest evidence that clonal phenotype is TCR-encoded came from comparing clones across donors. The HLA-A*02-restricted NLVPMVATV response contains public TCR motifs^43^, and our TCRdist analysis recovered both public alpha and beta chains across our donors. We identified major clones in seven TCR clusters and found aa-identical TCRs shared across donors in four out of the five pp65-specific clusters. We further found that clones within a TCR cluster occupied shared similar transcriptional neighborhoods, supporting a link between TCR sequence and functional specification. That unrelated individuals independently generate near-identical receptors, and that those receptors then give rise to similar differentiation states, is a clear indication that TCR sequences, presumably through a combination of avidity, docking geometry and the quality of downstream signaling, influence cell differentiation and transcriptional phenotype.

Consistent with prior studies^27,28^, we observed a remarkable degree of clonal stability during long periods of viral latency. Across all five donors examined, 47 out of 48 major clones we identified were detected at both timepoints, clonal hierarchies were highly preserved in four of five donors, and the fraction of a clone’s cells falling into the Tcm, Tem and T-CMC1 phenotypes were highly correlated across intervals of 4.5-12.5 years. Thus, we show that stability holds both at the level of clonal dominance and clonal phenotype, over intervals exceeding a decade. We observed no net expansion of either specificity across these intervals, and no correlation between the size of the CMV-specific pool and donor age in our broader screening cohort, which is difficult to reconcile with models of continuous memory inflation in humans^6,44,45^. However, these models are largely based on studies in MCMV infection in mice or on cross-sectional studies of single samples taken from younger or older cohorts and therefore do not monitor the dynamics of CMV-specific CD8^+^ T cells over time within individuals. Our results suggest that memory inflation in humans is not a constant process, but rather that the size of the CMV-specific T cell pool either increases episodically due to extrinsic factors which we did not observe in our latent cohort, or that memory inflation primarily occurs after other late-age-associated immune changes.

Several limitations of this study should be noted. Our longitudinal cohort comprised five donors with two timepoints each, so although the internal comparisons are well controlled, our power was limited to assess how these patterns vary across donors with age, sex, disease status or other confounding factors. The cohort also included both healthy control donors and donors with type 1 diabetes who were not receiving immune-modifying therapies, and we were not powered to test disease status as a variable. We sampled high-frequency responses from peripheral blood only, and clones with lower frequencies or tissue-restricted distributions may be systematically under-represented. Our analysis was also restricted to two viral antigens presented by three HLA class I alleles. Although the pp65-vs-IE1 distinction was reproducible across these alleles, we cannot say whether a comparable divergence would apply to the many other CMV antigens against which responses are mounted, or to HLA-B, HLA-C or HLA-E-restricted responses, which may also be important contributors to the response^46,47^. Within each of our donor’s non-HLA-A*01/02/24 alleles, there are likely many other CMV-specific T cell pools which may impact how the responses we monitored developed. For our acute infection analysis, we had data from only a single individual, so the timing we infer for the acute-to-latent transition requires confirmation in additional subjects. Finally, the relationship between relative TCR avidity/TCR sequence and transcriptional phenotype in this study is only correlative, although the recurrence of the same TCR-phenotype pairing across unrelated donors strengthens our conclusions.

Our findings carry practical implications for efforts to harness CMV-specific T cells therapeutically. Strategies that re-target CMV-specific T cells against tumors generally treat an epitope-specific population as functionally uniform. Our data indicate that it is not, and that the relevant functional unit is the clone. The observation that similar TCRs specify convergent fates further suggests that TCR sequence itself could serve as a selection criterion for identifying clones that reproducibly generate highly proliferative, exhaustion-resistant progeny, to specifically re-target these clones against tumors or use these clones as platforms for CARs. The same logic applies to CMV-based vaccine strategies, where the choice of which endogenous epitope to replace with a vaccine antigen may determine not only the magnitude of the resulting response but its differentiation state and durability. More broadly, CMV infection constitutes a natural setting in which the same T cells are stimulated repeatedly for decades without functional collapse. That individual clones can hold stable and distinct differentiation states throughout this process suggests that durable, functional heterogeneity in CD8^+^ T cells is possible even in the face of chronic stimulation, a state which is generally thought to cause T cell senescence and exhaustion.

## Methods

### Human participants

The Sound Life Project was approved by BRI’s Institutional Review Board (IRB19045). Samples from people with diabetes and healthy subjects were collected as part of BRI’s Immune-Mediated Disease Registry and Repository and were approved under protocols IRB10024 and IRB3041700, respectively. All subjects provided written informed consent in accordance with the Declaration of Helsinki.

### Acutely infected donor

Cryopreserved PBMC from the acutely infected donor were obtained from a single healthy participant enrolled in the Sound Life Project, a longitudinal study of peripheral blood immune phenotype in healthy adults in the Puget Sound region (ref, Smithmyer et al.). During the study monitoring period this participant seroconverted from CMV-seronegative to CMV-seropositive within a 4.5-month window between blood donations. Samples from five serial visits were analyzed, comprising one pre-seroconversion visit and four post-seroconversion visits spanning 16 months in total.

### Clinical laboratory PBMC/serum isolation, testing, and cryopreservation

Venous blood from study volunteers was collected in vacutainer tubes containing spray-coated silica (to prevent red cells from sticking to the tube wall) and a polymer gel for serum separation, then spun at 1400xg for 20 min. The cellular fraction was resuspended in phosphate buffered saline (PBS) and PBMC were separated from red blood cells using Ficoll extraction and frozen at -80°C before being stored in liquid nitrogen. PBMCs were thawed at 37°C and washed twice before use. HCMV serology testing was performed at the University of Washington’s Clinical Virology Laboratory in the Department of Laboratory Medicine using the Food and Drug Administration–approved LIAISON CMV IgG assay to qualitatively detect CMV IgG class antibodies. Results were reported as positive or negative for each sample.

### Peptide–MHC class I tetramer generation

Peptide–MHC class I tetramers were used to identify CD8+ T cells specific for pp65 and IE1. For the acutely infected donor, tetramers presented the pp65 peptide YSEHPTFTSQY on HLA-A01 and the IE1 peptide AYAQKIFKIL on HLA-A24. For the latently infected cohort, tetramers presented the pp65 peptide NLVPMVATV and the IE1 peptide VLEETSVML on HLA-A*02. The HLA-A*02 pp65 tetramer was obtained from the NIH tetramer lab, all other tetramers were synthesized in house using UV cleavable MHC-1 monomers. For these tetramers, peptides were synthesized at minimum 85% purity and provided lyophilized by Mimotopes (Australia) or GenScript (USA). Peptides were resuspended to 10mM stock concentration in DMSO and diluted to 1mM for experimental usage. UV cleavable HLA monomers were produced by the Immune Monitoring Core at Fred Hutchinson Cancer Center (HLA-A*01- STAPGJLEY, HLA-A*02 - GILGFVFJL, HLA-A*24-AYIDNJKF; “J” indicates UV labile amino acid). The peptide of interest was mixed with its cognate HLA in a 96 well U bottom plate, placed on ice and exposed to 368nm UV radiation (UV Crosslinker, Fisher Scientific cat # 13-245-221) for two 5 minute incubations, rotating the plate 180° after the first incubation. Exchanged peptide-MHC I complexes were incubated overnight at 4°C. Tetramers were produced by adding fluorescent conjugated streptavidin to the peptide-MHC complexes at a 1:4 ratio at 4 timed intervals (5 minutes x 3 followed by 10 minutes x 1), at room temperature.

### Flow cytometry screening of PBMCs

For surface phenotyping, 5-20x10e6 PBMC per sample were divided into four 5mL polystyrene tubes and cells were pelleted at 250 x g for 5 minutes. Pellets were resuspended at 5x10e6/mL in one of the following treatment conditions: DMSO (Sigma-Aldrich, >99.5% cell culture grade), or purified pp65 or IE1 peptide at 200 ng/ml. Stimulation was performed for 18 hours in RPMI media supplemented with 10% FBS, 1% glutamax, 1% penicillin/streptomycin, and 0.05% βME. After stimulation, cells were stained with pp65 and IE1 tetramers, as well as surface antibodies for T cell phenotype and activation (antibodies listed in reagents table).

### Flow cytometry

Data were acquired on a five-laser Cytek Aurora. Control PBMCs or UltraComp eBeads (ThermoFisher) were used for compensation. Data were analyzed using SpectroFlow (Cytek Biosciences) and FlowJo10 (Becton Dickinson) software.

### Cell sorting for single-cell sequencing

For single-cell sequencing, cells were sorted by fluorescence-activated cell sorting. CD8+ T cells were first enriched from one vial per timepoint by negative magnetic selection, and pp65-tetramer-positive, IE1-tetramer-positive, and non-tetramer-binding bulk CD8+ T cells were sorted from each of two blood draws per donor, or each of four visits for the seroconverting donor. (Fig S4A). Cells were sorted on a five-laser BD FACSAria Fusion using a 100-micron sort nozzle.

### CITE-seq cell hashing

For single-cell library preparation, sorted populations were labeled with an oligonucleotide-conjugated (CITE-seq) antibody panel including cell-hashing antibodies, then pooled prior to droplet encapsulation. As indicated in Fig S4A for the latently infected cohort, cells from each timepoint were stained with the sorting and CITE-seq panels, split into three fractions (approximately 10% / 45% / 45%), each labeled with one of six hashing barcodes; the corresponding fractions from the two timepoints were then pooled and sorted into tetramer-negative, pp65-tetramer-positive and IE1-tetramer-positive populations for encapsulation. In the analysis, hashtag identity encoded both the sorted timepoint and the antigen specificity of each population. Donors 4 and 5 were processed together as a multiplexed pool requiring hashtag demultiplexing, whereas donors 1 to 3 were not, which produced differing cell-barcode formats between the two batches. For the seroconverting donor, visits 1 and 2 were processed together, and visits 3 and 4 were processed together.

### Single-cell library preparation and sequencing

A single cell suspension was prepared from sorted cells and loaded onto the 10x Chromium Controller (10X Genomics) according to the manufacturer’s protocol. Sequencing libraries for gene expression, TCR, and feature barcoding were generated using the NextGEM Single Cell 5’ v2, or GEM-X Single Cell 5’ Reagent v3 kits (10x Genomics). Libraries were pooled and quantified using a Qubit® Fluorometer (Life Technologies). Sequencing of pooled libraries was carried out on a NextSeq 2000 sequencer (Illumina), with a target depth of 20,000 reads/cell for GEX and 5,000 reads/cell each for TCR and feature barcoding libraries. Data processing and quality control metrics were generated using CellRanger (10x Genomics).

### Bulk TCR sequencing

Bulk TCR sequencing of the sorted fractions was performed using the SMARTer Human TCR a/b Profiling Kit (Takara Bio) to prepare libraries, which incorporates unique molecular identifiers (UMIs) for template counting and error correction. Libraries were pooled equimolarly and quantified using a Qubit® Fluorometer (Life Technologies). Sequencing of pooled libraries was carried out on a NextSeq 2000 sequencer (Illumina) with paired-end 301-base reads, using a NextSeq™ 2000 XLEAP-SBS™ sequencing kit (Illumina). Base calls were processed to FASTQs on BaseSpace (Illumina), and a base call quality-trimming step was applied to remove low-confidence base calls from the ends of reads. The FASTQs were aligned to the GRCh38 human reference genome, using STAR v.2.4.2a and gene counts were generated using htseq-count. QC and metrics analysis was performed using the Picard family of tools (v1.134).

### Publicly available dataset

The pan-cancer CD8+ T cell atlas of Zheng et al. (2021) was obtained from Gene Expression Omnibus under accession GSE156728. Per-cancer-type count matrices were downloaded, loaded into Seurat, and merged across cancer types for gene-set scoring.

### Single-cell data preprocessing

Sequencing reads were processed with Cell Ranger (10x Genomics), using Cell Ranger for demultiplexing and alignment against the human reference transcriptome, with paired TCR contigs assembled by the V(D)J pipeline and libraries aggregated across sequencing runs. Hashtag identities were assigned to cells by thresholding normalized (log-CPM) hashtag-barcode counts, and doublets were removed. Cells were retained that had between 250 and 6,000 detected genes and between 450 and 35,000 UMIs, no more than 12.5% of reads mapping to mitochondrial genes, no more than 10% mapping to hemoglobin genes, and between 7.5% and 50% mapping to ribosomal genes.

### Integration, clustering and dimensionality reduction

Gene expression data were log-normalized, and the 2,000 most variable genes were identified by the variance-stabilizing transformation. To integrate cells across donors, each dataset was split by donor, normalized and scaled independently, and integrated using Seurat’s canonical correlation analysis anchor-based. Principal component analysis was performed on the scaled data, and the first 30 principal components were used for Uniform Manifold Approximation and Projection (UMAP) and for shared-nearest-neighbor graph construction, followed by Louvain community cluster detection. For the acutely infected donor, this yielded 8 transcriptional clusters. For the latently infected cohort, an initial clustering that included tetramer-negative bulk CD8+ T cells was followed by subsetting to tetramer-binding cells only, which were re-clustered into 9 clusters.

### Clonotype definition

Cells were assigned to clonotypes on the basis of shared TCRα and TCRβ nucleotide sequences using the tcrGraph package (BenaroyaResearch/tcrGraph), operating on productive, high-confidence contigs. For the acutely infected donor, major clones were defined as clonotypes comprising more than 1% of all sampled tetramer-binding cells of a given specificity across all visits combined. For the latently infected cohort, major clones were defined as clonotypes with at least 20 cells detected across both timepoints combined, a threshold set to ensure adequate power to estimate each clone’s phenotype composition. Clonal identity was defined at the nucleotide level throughout, so that clones sharing an identical CDR3 amino acid sequence but differing in nucleotide sequence were treated as distinct.

### Clonal diversity and stability

Clonal repertoire diversity was quantified using the Gini–Simpson index (1 − Σpᵢ², the probability that two randomly chosen cells belong to different clonotypes). For the acutely infected donor, per-visit Gini–Simpson indices were reported with bootstrap 95% confidence intervals (R = 2,000 resamples), and pairwise differences between visits were tested by permutation on the absolute difference in Gini–Simpson index (R = 10,000 permutations), with Holm-adjusted p values within each specificity (Fig S1G). Stability of each clone’s transcriptional distribution over time was quantified using the Jensen–Shannon divergence (computed with base-2 logarithms) between the clone’s distribution across clusters at successive timepoints; for the acute donor, phenotypic stability relative to the first visit was expressed as 1 − Jensen–Shannon divergence, computed for clones with at least 10 cells present at the visit in question (Fig S2C). Stability of clonal dominance across the two timepoints in the latently infected cohort was assessed by correlating each major clone’s frequency (as a percentage of its antigen-specific pool) at the early versus the late timepoint, with a linear regression reported (R² and slope p value; Fig 3C).

### Clonal phenotype stability

Clonal phenotype stability was assessed for all clones with at least 20 cells at both timepoints by calculating the percentage of each clone’s cells falling into each phenotype (Tcm, Tem, NK-like, ISG+, T-CMC1) at each timepoint, and correlating the early-timepoint and late-timepoint percentages for each phenotype by linear regression (Fig 3D–F, Fig S8B–C). The Gini–Simpson index was also applied across a clone’s own phenotype composition to quantify within-clone phenotypic heterogeneity (Fig 2K). Phenotype percentages were compared between pp65- and IE1-specific cells across the five donors by paired, two-tailed t test performed per phenotype (Fig 2F).

### TCR sequence clustering

Clusters of T cells with convergent TCR sequences were identified using TCRdist-based sequence similarity clustering of paired TCRα and TCRβ chains including CDR3, V and J gene regions. Conservative distance thresholds were used; 64 for the latently infected cohort inclusive of 5 people (Fig 5), and a more conservative threshold of 48 was used for the acutely infected donor (as this was performed within-person). These thresholds reflect up to 2 non-synonymous amino acid changes per TRA/TRB CDR3 at a TCR distance of 48 and 2 or 3 per chain at 64.

### kNN phenotypic bias metric

Phenotypic coherence of the TCR distance clusters was quantified with a k-nearest-neighbor bias metric. For each cell, the 100 nearest neighbors were identified in the 30-dimensional integrated principal-component space after excluding neighbors belonging to the same clonotype and restricting candidate neighbors to cells of the same antigen specificity (each TCR cluster was assigned a single specificity by majority vote of its cells). For each cluster, the metric was the number of a cell’s neighbors belonging to the same TCR cluster, averaged first within each clone and then across clones so that abundant clones could not dominate the score. Significance was assessed against a null distribution generated by permuting cluster labels within specificity (10,000 permutations); the empirical p value was (1 + number of null values ≥ observed) / (permutations + 1), and p values were adjusted across clusters by the Benjamini–Hochberg procedure. The reported effect for each cluster was its observed score minus the mean of its null distribution (Fig 5D). Excluding same-clone neighbors ensures that the metric reflects convergence of transcriptional state across distinct clones sharing a TCR cluster, rather than the internal coherence of individual clones.

### Bulk TCRseq enrichment scores

For TCR avidity and proliferative capacity scores, TCRβ clonotypes were extracted from each bulk TCR-sequencing library using TRUST4. Bulk clonotypes were matched to the single-cell-derived clonotypes by TCRβ CDR3 nucleotide sequence. Clones supported by at least one UMI were retained, and within each sorted well a per-clone UMI count and frequency were computed. Clone abundances were then normalized within each well by the centered log-ratio transformation, with a pseudocount of 1 added before transformation. A relative TCR avidity score was defined for each clone as its centered-log-ratio abundance in the tetramer-high sort minus its abundance in the tetramer-low sort, and a proliferation score as its centered-log-ratio abundance in the proliferated fraction minus its abundance in the unsorted control (Fig 4B–C, Fig S9). Correlations of these scores with one another, with clonal frequency, and with phenotype composition were assessed by linear regression (Fig 4D–J, Fig S9D–F), and technical reproducibility was evaluated by comparing scores between independent assay replicates (Fig S9A, S9C).

### Statistics

Statistical analyses were performed in R unless otherwise noted, with statistical details for each experiment — the test used, the value of n, and what n represents — given in the corresponding figure legend. Comparisons between paired antigen-specific populations within donors used paired t tests; permutation and bootstrap procedures for diversity metrics are described above; and correlations were assessed by linear regression with R² and slope p values reported, or, where indicated, by Spearman rank correlation with Benjamini–Hochberg correction.

### Software

Analyses used R v4.5.1, Seurat v 5.3.0, tcrGraph, UCell, the compositions, boot, philentropy and FNN R packages, Cell Ranger v10, FlowJo v10, GraphPad Prism 10. Cluster analysis was performed using python v3.12.12, tcrdist3, tcrdistgpu.

### Data and code availability

Single-cell RNA-sequencing, CITE-seq and paired TCR-sequencing data, together with the bulk TCR-sequencing data, are available at dbGaP and GEO. All original analysis code has been deposited at github.com/BenaroyaResearch.

## Supporting information

Key Resources Table

## Acknowledgements

We thank Rachel Hartley and Thien-Son Nguyen for assisting with sample and clinical data tracking; the clinical coordinators and technicians in the Center for Interventional Immunology for sample collection and processing; Dr. Adam Wojno, Dr. Amir Figueroa and the BRI Cell and Tissue Analysis Core for flow cytometry support. Collection of samples and data for the Sound Life Project was supported by the Allen Institute for Immunology and by philanthropic donations to BRI through the Virginia Mason Health Foundation. This work was supported by the Leonard and Norma Klorfine Foundation. The graphical abstract was created with BioRender.com.

## Supplemental Figure Legends

**Figure S1.**
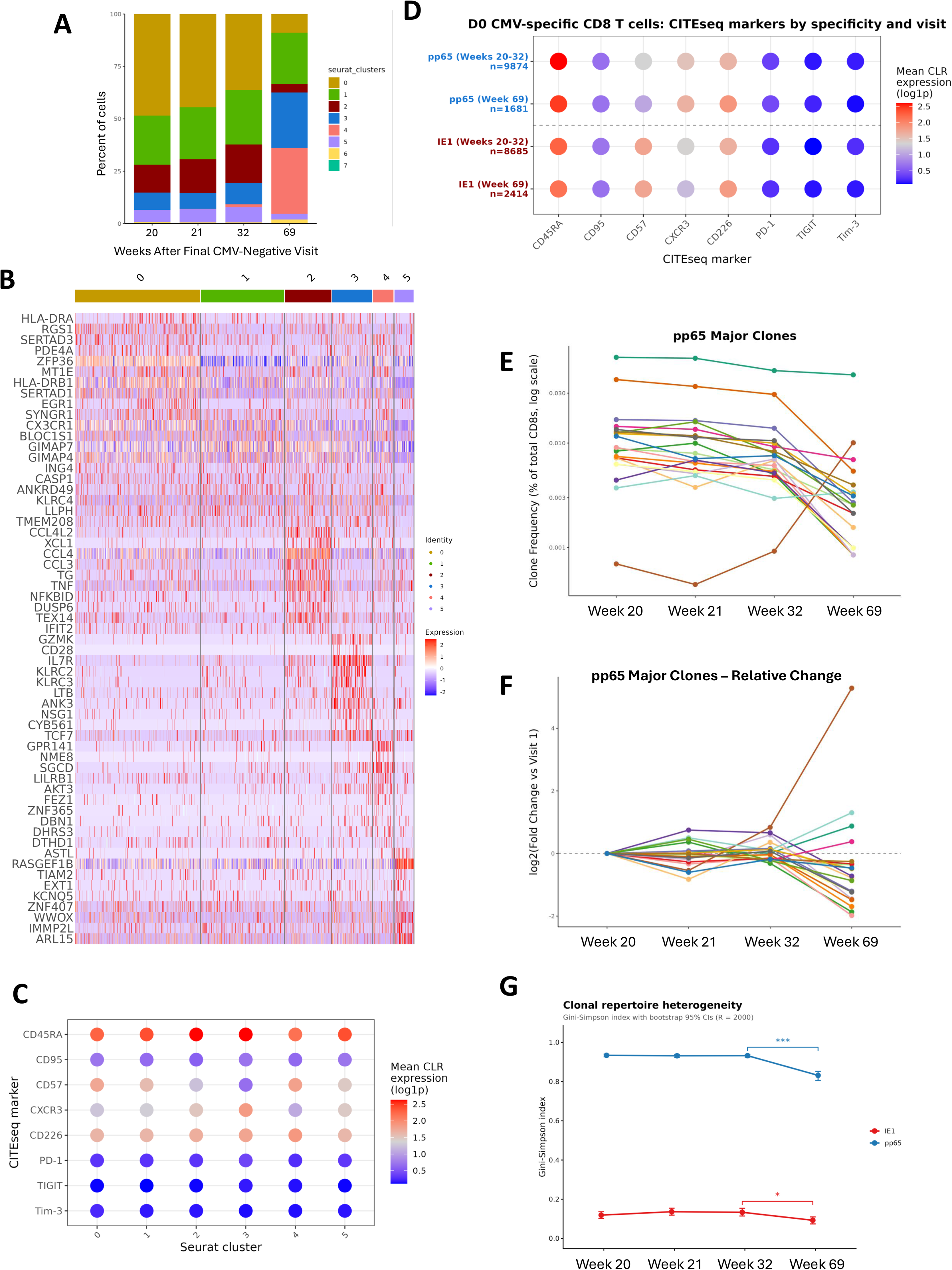
Phenotypic and clonal characterization of CMV-specific T cells during early infection. Related to Figure 1. **A.** Stacked bar plot of the percentage of all CMV-specific CD8^+^ T cells assigned to each Seurat cluster at each visit. **B.** Heatmap of the top differentially expressed genes defining the main Seurat clusters 0–5. Color indicates scaled expression. **C.** Dot plot of CITEseq surface markers across Seurat clusters. Dot color indicates mean centered log ratio (CLR) normalized marker expression. **D.** Dot plot of the same CITEseq markers, split by specificity and visit. Dot color indicates mean CLR-normalized marker expression. **E.** Frequency of each pp65-specific major clone across weeks 20-69, expressed as a percentage of total CD8^+^ T cells, on a log scale. **F.** Relative change in frequency of each pp65-specific major clone across visits, expressed as log2 fold change vs week 20. **G.** Gini-Simpson index of clonal diversity (repertoire heterogeneity) for the pp65- and IE1-specific pools at each visit, with bootstrap 95% confidence intervals (R = 2,000). Brackets show pairwise permutation tests on |ΔGini-Simpson| (R = 10,000), Holm-adjusted p-values within specificity. *p < 0.05, **p < 0.01, ***p < 0.001.

**Figure S2.**
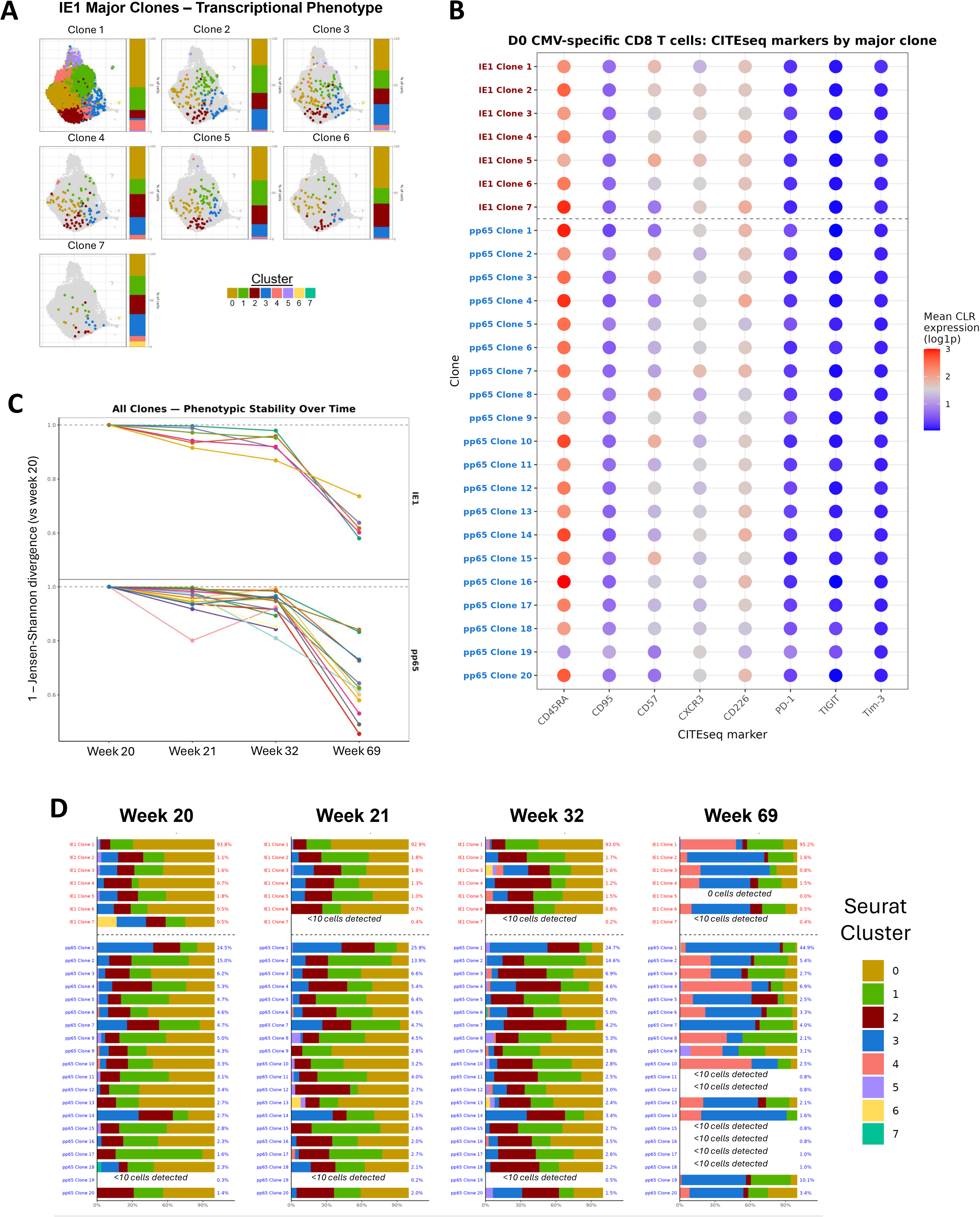
Clonal heterogeneity and differentiation trajectories during early infection. Related to Figure 1. **A.** UMAP position of each IE1-specific major clone. Each panel shows cells belonging to that clone colored by Seurat cluster; the adjacent stacked bar graph shows the cluster composition percentages. **B.** Dot plot of CITEseq marker expression for each major clone. Dot color indicates mean CLR-normalized marker expression. **C.** Phenotypic stability of each clone over time, plotted as 1 − Jensen-Shannon divergence between the clone’s distribution across the Seurat clusters at a given visit and its distribution at week 20. Higher values indicate greater similarity to week 20. Clones plotted for all visits with at least 10 cells present at that visit. **D.** Stacked bars of Seurat cluster composition for each major clone at each visit, with the clone’s percentage of its antigen-specific pool given to the right of each bar.

**Figure S3.**
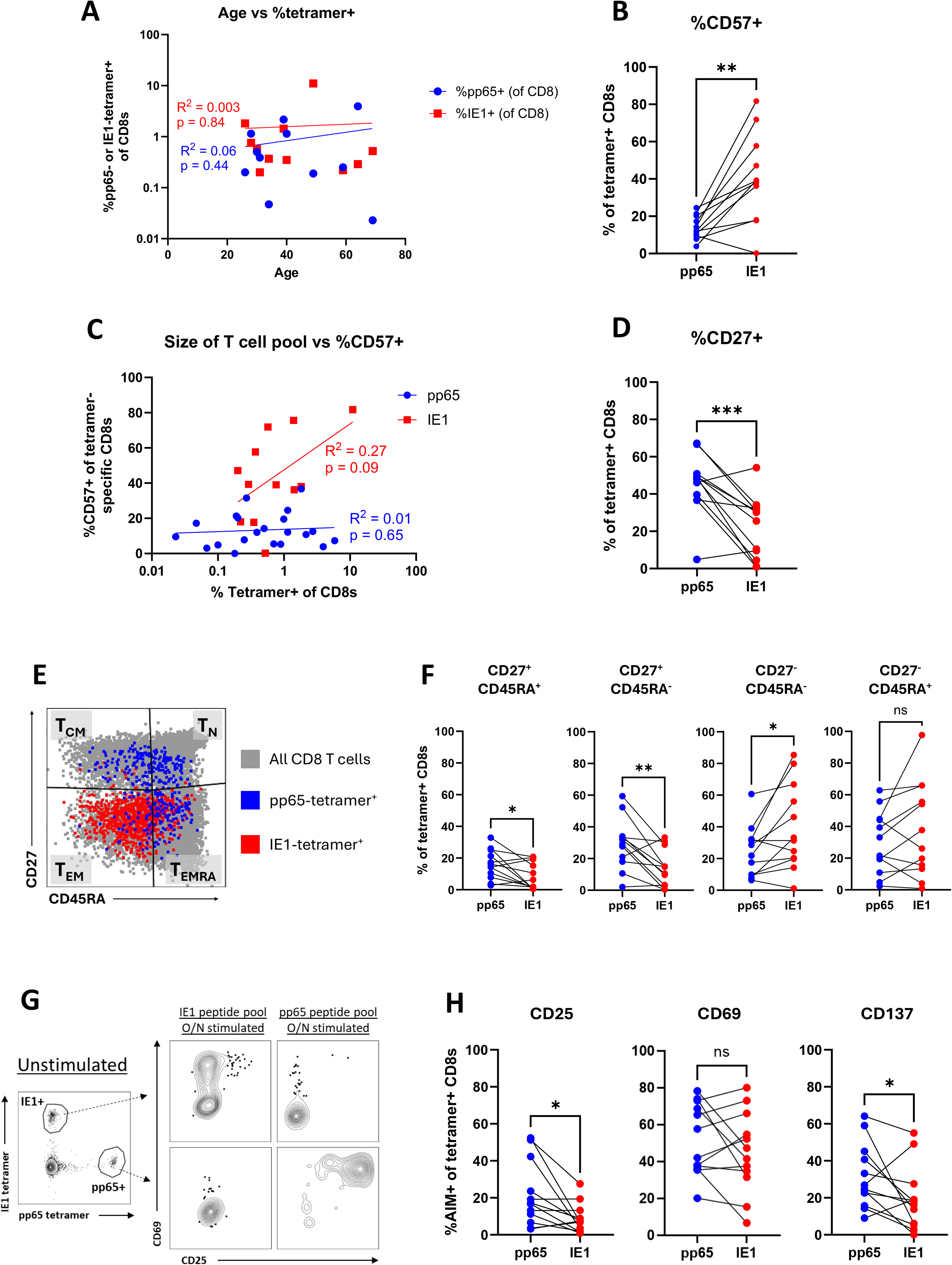
Flow cytometric phenotyping of pp65- and IE1-specific T cells in the latently infected cohort. Related to Figure 2. **A.** Donor age vs the frequency of pp65- and IE1-tetramer+ cells as a percentage of CD8^+^ T cells (log scale), with linear regression shown per specificity. **B.** Percentage of CD57 expression among pp65- and IE1-tetramer+ CD8^+^ T cells, paired by donor. **C.** Size of the antigen-specific T cell pool (% tetramer+ of CD8s, log scale) vs the percentage of CD57 expression within that pool, with linear regression shown per specificity. **D.** Percentage of CD27 expression among pp65- and IE1-tetramer+ CD8^+^ T cells, paired by donor. **E.** Representative CD27 vs CD45RA plot with Tn-, Tcm-, Tem- and Temra-like quadrants, overlaying pp65- and IE1-tetramer+ cells onto all CD8^+^ T cells. **F.** Frequency of each CD27/CD45RA subset among pp65- and IE1-tetramer+ CD8^+^ T cells, paired by donor. **G.** Representative activation-induced marker (AIM) assay. Left, unstimulated sample showing the HLA-A*02:IE1 tetramer and HLA-A*02:pp65 tetramer gates. Right, CD69 vs CD25 expression on the gated tetramer+ populations after overnight stimulation with the IE1 or pp65 peptide pool. **H.** Percentage of AIM+ cells among pp65- and IE1-tetramer+ CD8^+^ T cells after cognate peptide stimulation, paired by donor. *p < 0.05, **p < 0.01, ***p < 0.001.

**Figure S4.**
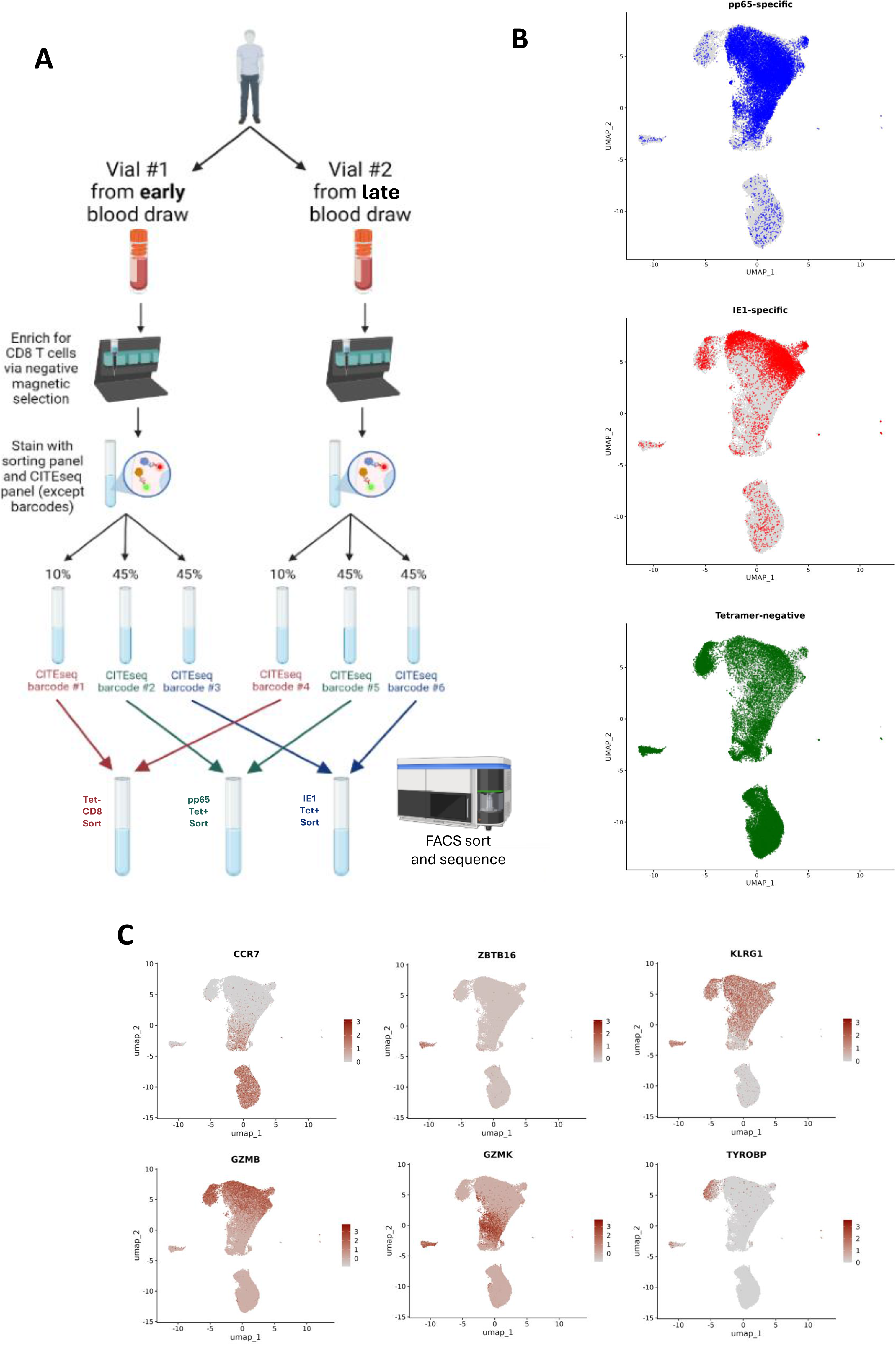
Experimental design and global transcriptional landscape of sorted CD8^+^ T cells from latent cohort. Related to Figure 2. **A.** Schematic of the sample processing workflow. For each donor, one vial from an early and one from a late blood draw were enriched for CD8^+^ T cells by negative magnetic selection, stained with the sorting and CITEseq panels (excluding hashing barcodes), then split 10%/45%/45% and labeled with one of six CITEseq barcodes. The corresponding tubes from each timepoint were pooled and FACS sorted for tetramer-negative CD8, pp65-tetramer+ and IE1-tetramer+ populations respectively, then sequenced. **B.** UMAP of integrated scRNAseq data from all sorted CD8^+^ T cells across all donors and timepoints, with pp65-specific, IE1-specific, and tetramer- negative cells highlighted on the same embedding. **C.** Feature plots of selected gene expression on the UMAP from (B). Color indicates normalized expression.

**Figure S5.**
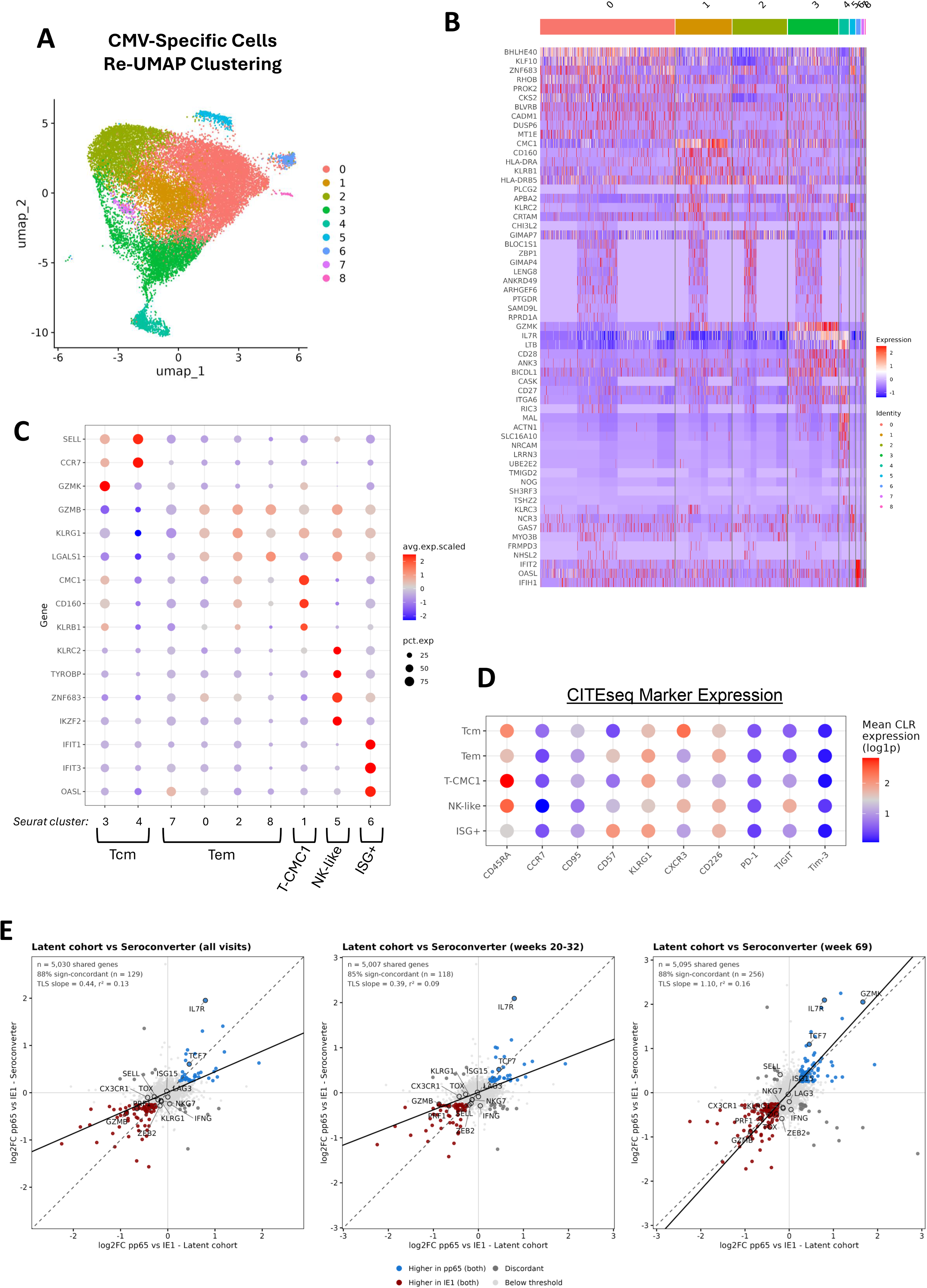
Definition of transcriptional phenotypes from CMV-specific cells of the latent cohort. Related to Figure 2. **A.** UMAP of re-clustered tetramer-binding CMV-specific cells only, colored by Seurat cluster. **B.** Heatmap of the top differentially expressed genes for each Seurat cluster. Color indicates scaled expression. **C.** Dot plot of selected marker genes across Seurat clusters, with brackets showing the assignment of clusters into phenotypes. **D.** Dot plot of CITEseq marker expression across the five phenotypes. **E** Concordance of pp65-vs-IE1 differential gene expression between the latently-infected cohort (x-axis) and the seroconverting donor (y-axis), for all of that donor’s visits (left), only weeks 20-32 (center) and only week 69 (right). Each point is a shared gene; genes significantly higher in pp65 in both datasets are blue, higher in IE1 in both are red, discordant genes grey, and genes below the significance threshold light grey. Solid black line, total least squares fit. Statistics are inset per panel.

**Figure S6.**
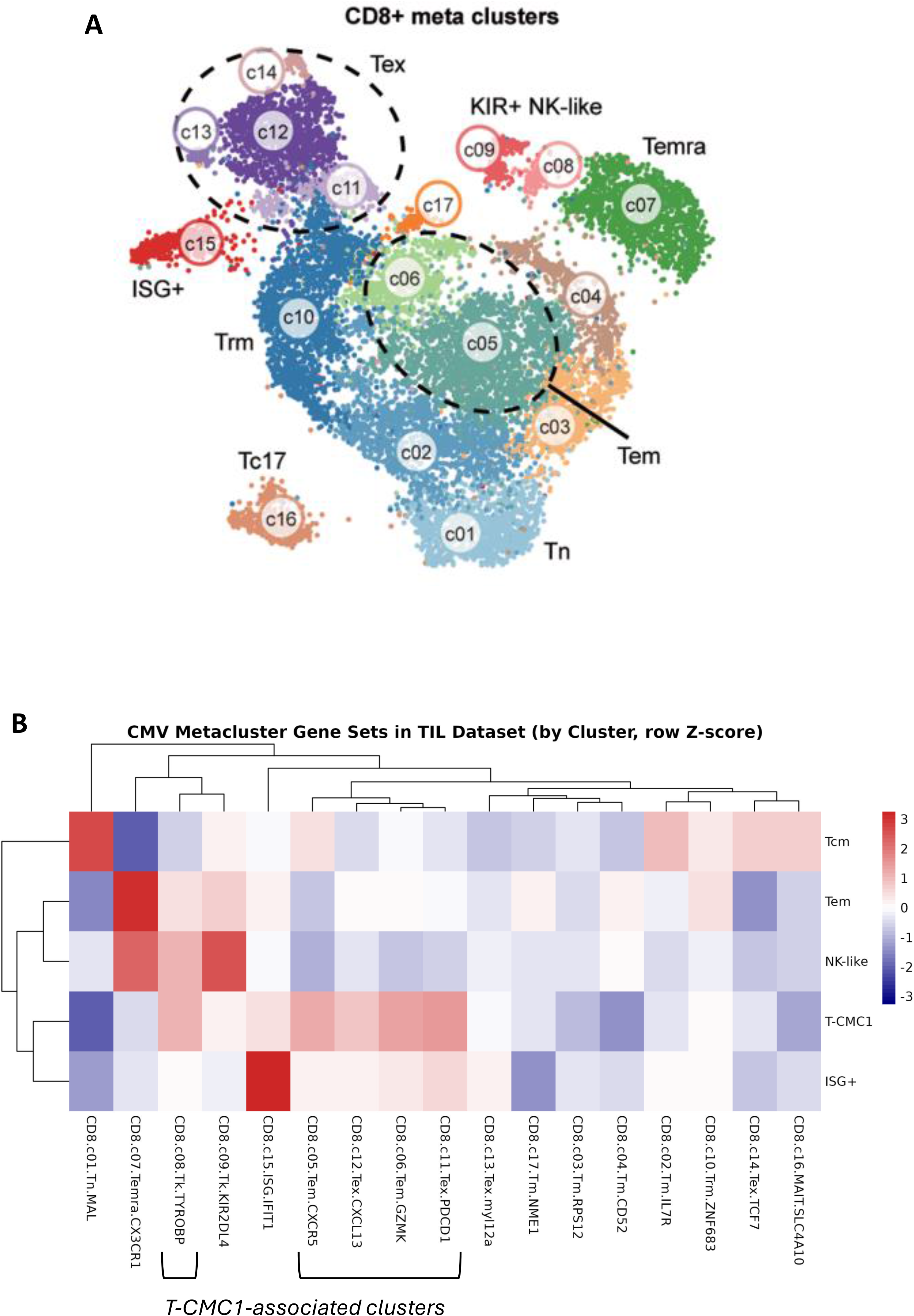
Comparison of CMV-specific phenotypes to a published pan-cancer CD8^+^ T cell atlas. Related to Figure 2. **A.** Reference UMAP of CD8^+^ T cell meta-clusters from the pan-cancer atlas (reproduced from reference 33), with the atlas’ 17 clusters grouped shown. **B.** Heatmap of gene set scores in the published atlas, where gene sets were generated from the top DEGs of each of our five phenotypes (rows) and scored across the atlas clusters (columns). Color indicates row Z-score; dendrograms show hierarchical clustering of rows and columns. Brackets mark the atlas clusters most strongly associated with the T-CMC1 gene set.

**Figure S7.**
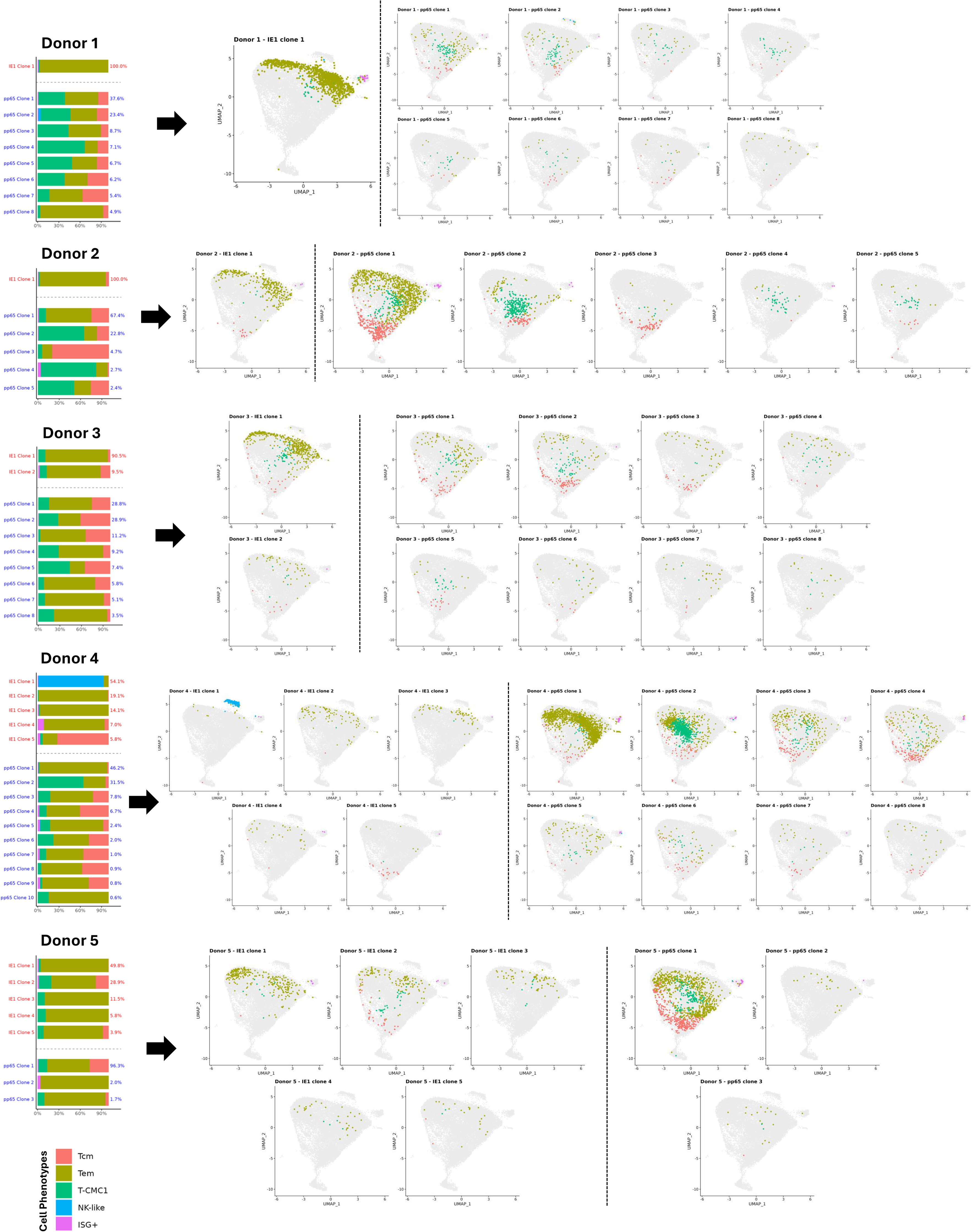
Clonally encoded transcriptional heterogeneity in each latently infected donor. Related to Figure 2. For each donor: left, stacked bars of phenotype composition for every major clone, with the clone’s percentage of its antigen-specific pool given to the right of each bar; right, UMAP position of each major clone, colored by phenotype, over a grey background of all CMV-specific cells from all donors.

**Figure S8.**
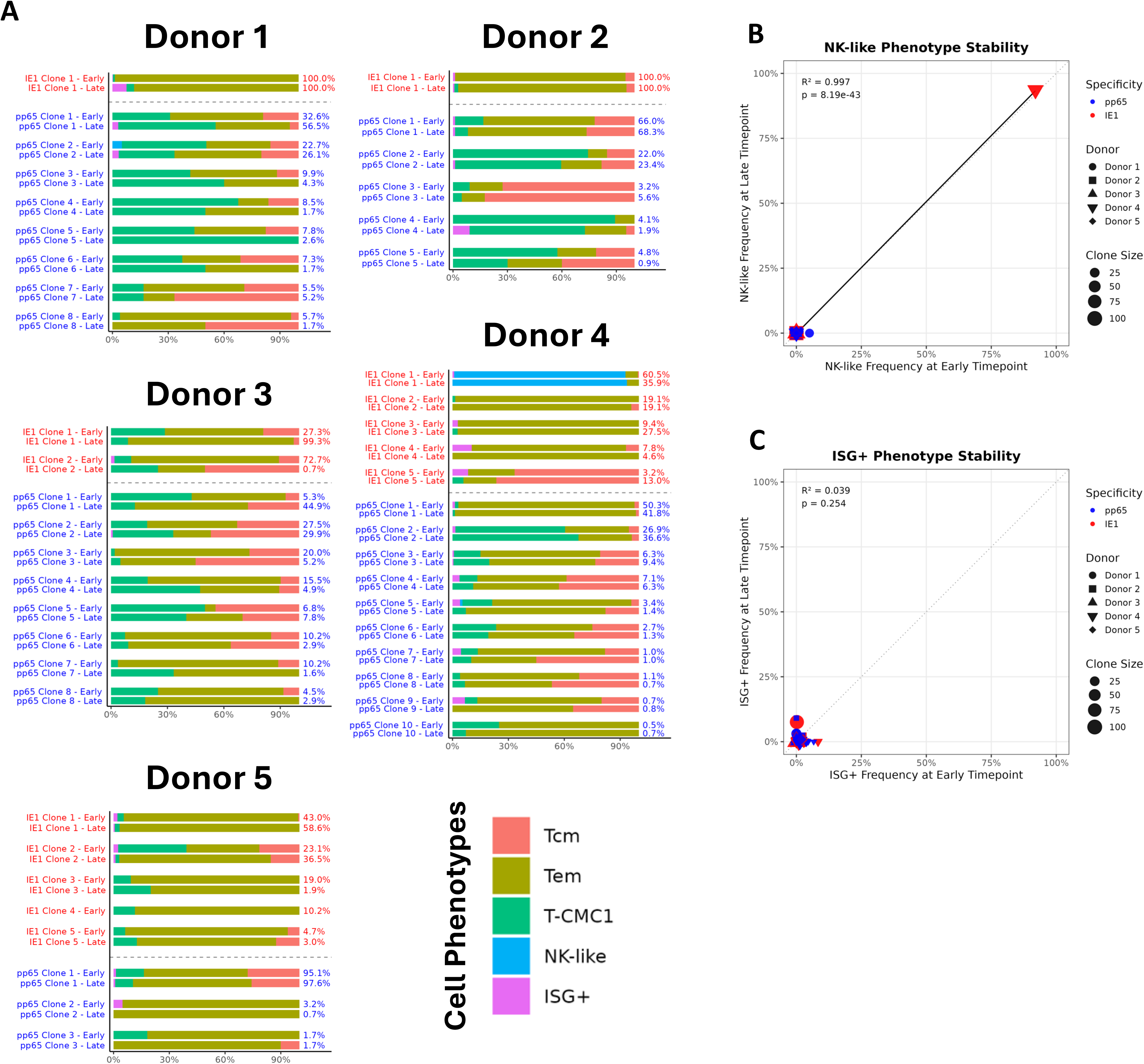
Stability of clonal transcriptional phenotype between timepoints. Related to Figure 3. **A.** For each donor, paired stacked bar plots of phenotype composition for every major clone at the early (top) and late (bottom) timepoint, with the clone’s percentage of its antigen-specific pool at each timepoint given to the right. Only clones with at least 20 cells are shown. **B.** Correlation between the percentage of cells within each clone assigned to the NK-like phenotype at the early vs the late timepoint. Points colored/shaped/sized as in Fig 3C–F. **C.** As in (B), for the ISG+ phenotype.

**Figure S9.**
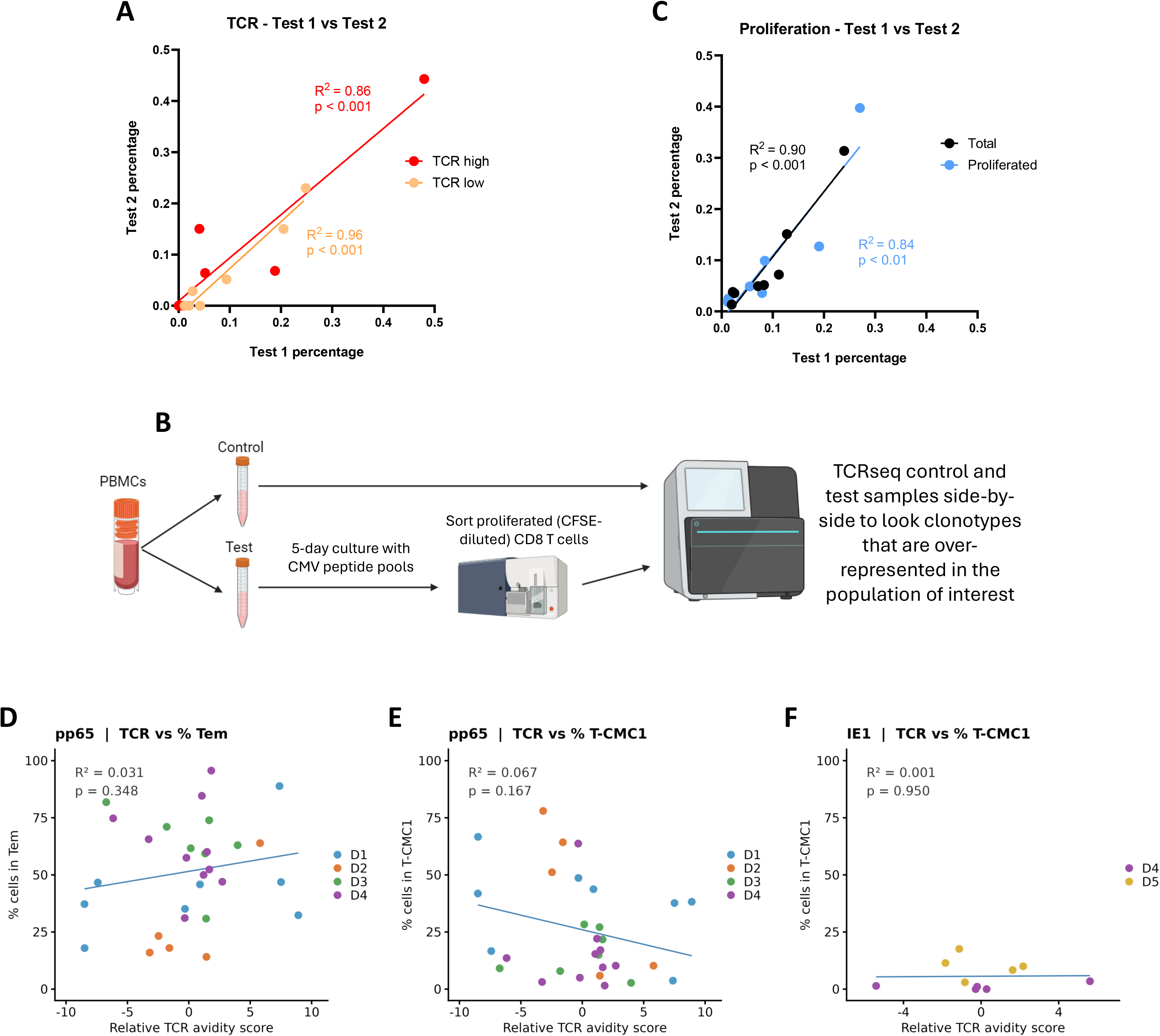
Bulk TCRseq enrichment assay design, technical reproducibility, and additional correlations. Related to Figure 4. **A.** Technical reproducibility of the tetramer high-vs-low MFI bulk TCRseq enrichment assay, shown as percent of each major pp65-specific clone from donor 1 detected in the TCR-high and TCR-low sorts, in repeat 1 vs repeat 2. Linear regression shown for the percentages in each gate. **B.** Schematic of the proliferation bulk TCRseq enrichment assay. PBMCs were split into a control and a test aliquot; the test aliquot was cultured with CMV peptide pools and IL-2 for 5 days, proliferated (CFSE-diluted) CD8^+^ T cells were sorted, and bulk TCRseq was performed on both aliquots in parallel to identify clonotypes over-represented in the proliferated population. **C.** Technical reproducibility of the proliferation bulk TCRseq enrichment assay, as in (A). **D.** Relative TCR avidity score vs percentage of cells with the Tem phenotype, pp65-specific clones. **E.** Relative TCR avidity score vs percentage of cells with the T-CMC1 phenotype, pp65-specific clones. **F.** Relative TCR avidity score vs percentage of cells with the T-CMC1 phenotype, IE1-specific clones.

**Figure S10.**
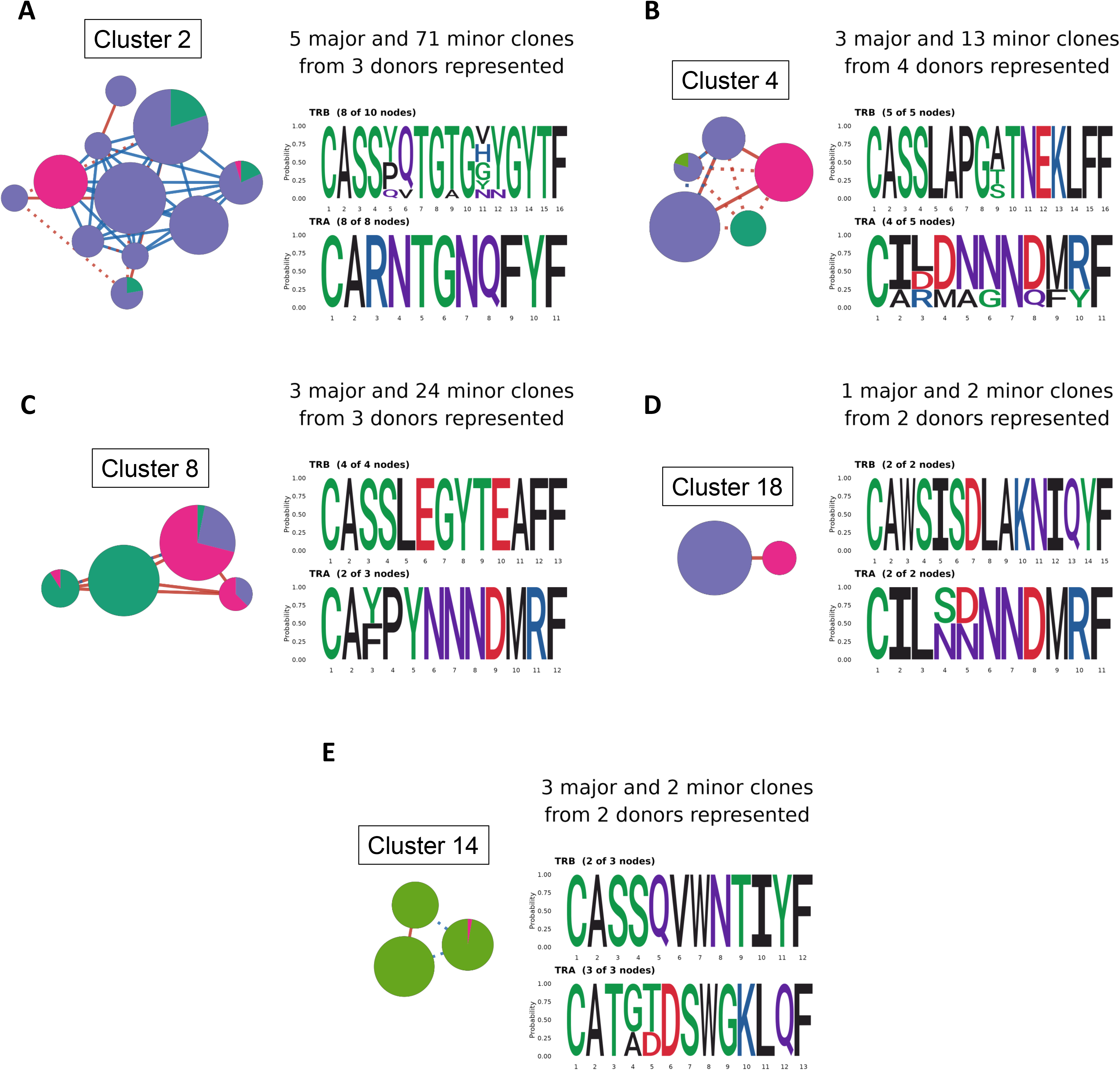
Logo-annotated constellation plots for the remaining convergent TCR clusters. Related to Figure 5. Constellation plots as in Fig 5B–C, with TRB (top) and TRA (bottom) CDR3 sequence logo for each node group and annotations giving the number of major and minor clones and contributing donors. **A.** pp65 Cluster 2. **B.** pp65 Cluster 4. **C.** pp65 Cluster 8. **D.** pp65 Cluster 18. **E.** IE1 Cluster 14.

**Figure S11.**
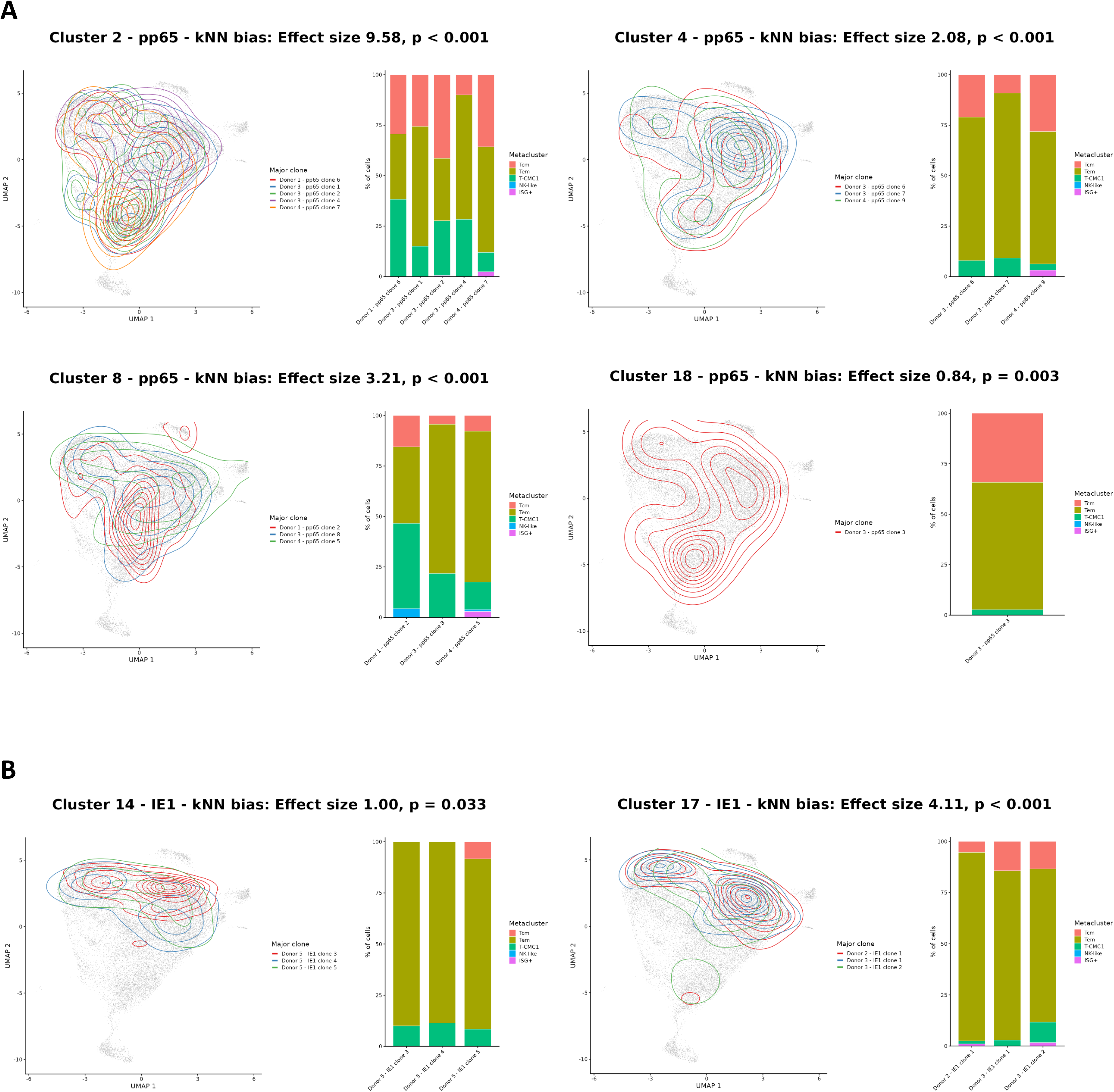
Shared transcriptional phenotypes among clones within convergent TCR clusters. Related to Figure 5. **A.** pp65-specific TCR clusters 2, 4, 8 and 18. For each cluster: kNN bias test effect size and significance over null hypothesis; UMAP density contours for each major clone over a grey background of all CMV-specific cells; and a stacked bar of phenotype composition for each major clone. **B.** As in (A), for IE1-specific TCR clusters 14 and 17.

**Table S1.**

| Donor | Disease Status | Sex | Race | pp65/IE1 Tetramer<br>HLA Restriction | Age at First Sequenced<br>Blood Draw | Age at Final Sequenced<br>Blood Draw | CMV Latency Verification |
| --- | --- | --- | --- | --- | --- | --- | --- |
| Seroconverter | Healthy | Male | White | A*01/A*24 | 36 | 38 | N/A |
| Latent Donor 1 | Healthy | Female | White | A*02 | 64 | 68 | CMV IgG+ at 2.5 years before first<br>draw |
| Latent Donor 2 | Type 1 Diabetes | Female | White | A*02 | 37 | 50 | CMV IgG+ at first draw |
| Latent Donor 3 | Type 1 Diabetes | Male | White | A*02 | 18 | 30 | pp65/IE1-tetramer binding cells found<br>at 1.5 years before first draw |
| Latent Donor 4 | Type 1 Diabetes | Male | White | A*02 | 23 | 29 | CMV IgG+/IgM- at first draw |
| Latent Donor 5 | Type 1 Diabetes | Male | White | A*02 | 17 | 26 | pp65/IE1-tetramer binding cells found<br>at 2 years before first draw |

## Notes

### Competing Interest Statement

The authors have declared no competing interest.

